# Umbrella toxin particles produced by Streptomyces block mycelial growth of competing species

**DOI:** 10.1101/2023.12.09.570830

**Authors:** Qinqin Zhao, Savannah Bertolli, Young-Jun Park, Yongjun Tan, Kevin J. Cutler, Pooja Srinivas, Kyle L. Asfahl, Citlali F. Garcia, Larry A. Gallagher, Yaqiao Li, Yaxi Wang, Devin Coleman-Derr, Frank DiMaio, Dapeng Zhang, S. Brook Peterson, David Veesler, Joseph D. Mougous

**Affiliations:** Department of Microbiology, University of Washington, Seattle, WA, USA; Howard Hughes Medical Institute, University of Washington, Seattle, WA, USA; Department of Biochemistry, University of Washington, Seattle, WA, USA; Department of Biology, Saint Louis University, St. Louis, MO, USA; Department of Physics, University of Washington, Seattle, WA, USA; Microbial Interactions and Microbiome Center, University of Washington, Seattle, WA, USA; Plant Gene Expression Center, USDA-ARS, Albany, CA USA; Department of Plant and Microbial Biology, University of California Berkeley, CA USA; Institute for Protein Design, University of Washington, Seattle, WA USA; Program of Bioinformatic and Computational Biology, St. Louis University, St. Louis, MO USA

**Author notes:** To whom correspondence should be addressed: Email –.

## Abstract

The *Streptomyces* are a genus of ubiquitous soil bacteria from which the majority of clinically utilized antibiotics derive. The production of these antibacterial molecules reflects the relentless competition *Streptomyces* engage in with other bacteria, including other *Streptomyces* species. Here we show that in addition to small molecule antibiotics, Streptomyces produce and secrete antibacterial protein complexes that feature a large, degenerate repeat-containing polymorphic toxin protein. A cryo-EM structure of these particles reveals an extended stalk topped by a ringed crown comprising the toxin repeats scaffolding five lectin-tipped spokes, leading to our naming them umbrella particles. *S. coelicolor* encodes three umbrella particles with distinct toxin and lectin composition, and supernatant containing these toxins specifically and potently inhibits the growth of select *Streptomyces* species from among a diverse collection of bacteria screened. For one target, *S. griseus*, we find inhibition relies on a single toxin and that intoxication manifests as rapid cessation of vegetative mycelial growth. Our data show that *Streptomyces* umbrella particles mediate competition between vegetative mycelia of related species, a function distinct from small molecule antibiotics, which are produced at the onset of reproductive growth and act broadly. Sequence analyses suggest this role of umbrella particles extends beyond *Streptomyces*, as we find umbrella loci in nearly one-thousand species across Actinobacteria.

## Introduction

Soil is typically home to a dense and diverse bacterial community, with many soils containing >10^9^ bacterial species per gram^1^. Under such conditions, interference competition is rampant, as evidenced by the wide array of interbacterial antagonism and defense systems these bacteria harbor^2,3^. The *Streptomyces* are a genus of ubiquitous soil bacteria that are notable for their production of antimicrobial secondary metabolites, many of which are used clinically as antibiotics^4–6^. Among other targets, *Streptomyces* spp. appear to use these antimicrobials to inhibit the growth of other *Streptomyces* spp., suggesting that interspecies antagonism within the genus is ecologically important^7^. In many bacteria, proteinaceous polymorphic toxins, in conjunction with their associated delivery machinery mediate interspecies competition^8–15^; however, such systems have not been identified in *Streptomyces*.

While polymorphic toxin delivery relies on distinctive, sequence divergent machineries particular to the producer and target species, the small toxin domains they transport often share homology. A comprehensive bioinformatic study exploiting this feature to search for new polymorphic toxins found that the uncharacterized alanine leucine phenylalanine-rich (ALF) repeat proteins of Streptomyces and related species bear C-terminal polymorphic toxin domains^14,16^. The model strain *S. coelicolor* encodes three ALF proteins, which we term umbrella toxin protein C 1-3 (UmbC1-3); each contains an N-terminal twin arginine translocation (TAT) signal, two sets of four ALF repeats (ALF1-8), two extended coiled-coil domains, and variable C-terminal and toxin domains (Figure 1a and Tables S1,S2). The ALF repeat is a degenerate (28% average identity across ALF repeats 1-8 from UmbC1-3) 43-44 amino acid motif of unknown function (Figure S1a)^16^.

**Figure 1:**
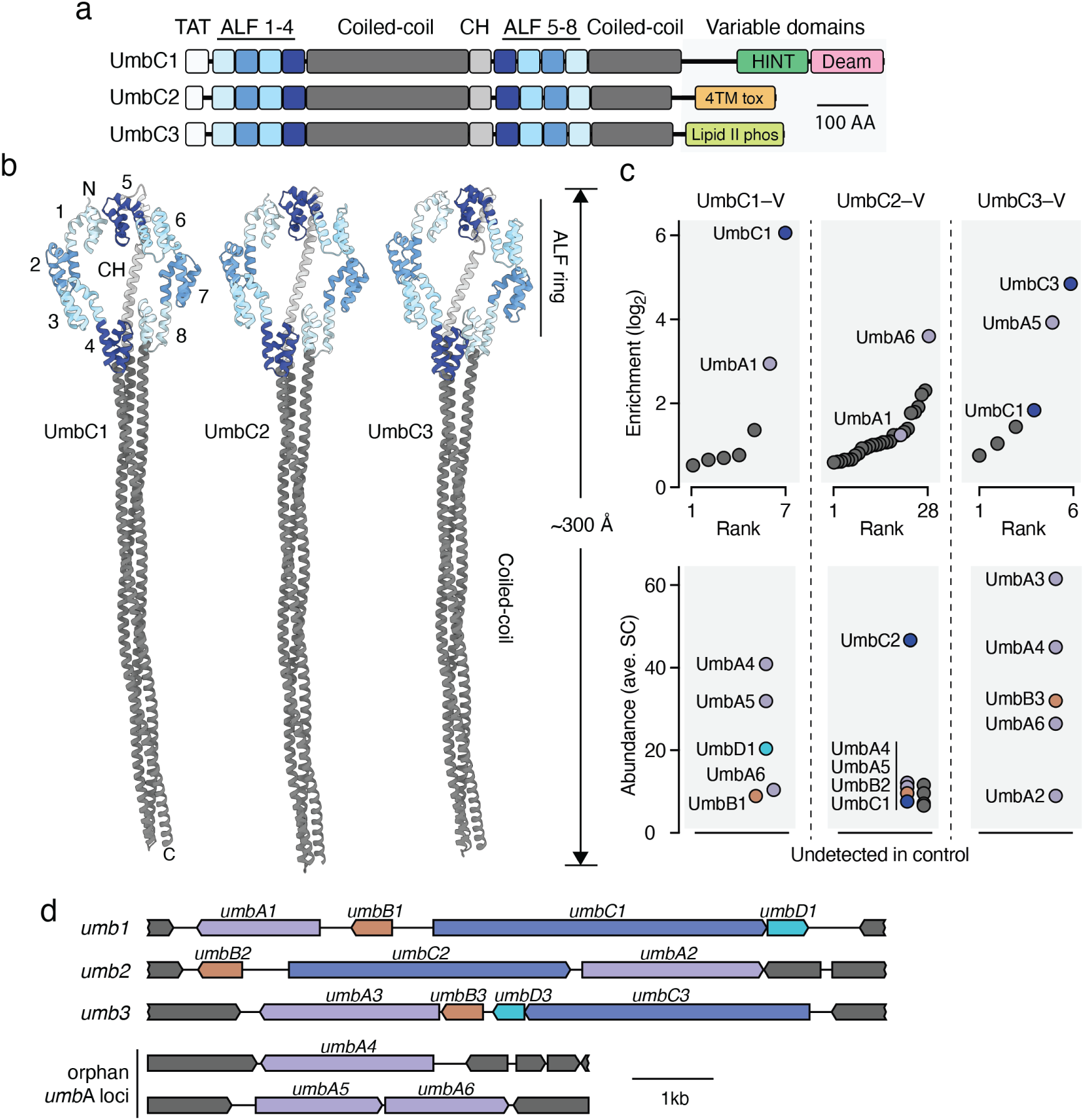
*S. coelicolor* encodes three degenerate repeat-containing polymorphic toxins which interact with paralogous proteins. **a,** Domain architecture of the UmbC proteins of *S. coelicolor*. TAT, twin-arginine translocation secretion signal; CH, connecting helix. Protein accession numbers and definition of variable C-terminal domains available in Tables S1 and S2. **b,** AlphaFold-predicted structural models of *S. coelicolor* UmbC proteins. UmbC1 and UmbC2 models were generated using template mode with UmbC3 as the reference. Colors correspond to (a); ALF repeat numbering and location of the connecting helix shown for UmbC1. The variable C-terminal domains, predicted to localize to the end of the stalk, could not be confidently modeled and thus are not shown. **c,** IP-MS identification of proteins that interact with UmbC1, UmbC2 or UmbC3 from *S. coelicolor.* Upper panels indicate the average fold enrichment of proteins detected in both IP and control samples; lower panels present abundance (average spectral counts, SC) for proteins detected only in IP samples. Colors indicate paralogous proteins; non-Umb proteins shown in grey. Note that additional background interacting proteins were identified for UmbC2, which we attribute to the lower abundance of this protein (46.5 SC) relative to UmbC1 (134.5 SC) and UmbC3 (781 SC) n = 2 biological replicates. **d**, Loci encoding Umb protein complex components in *S. coelicolor.* Orphan *umbA* loci are those encoded distantly from other complex constituents. Colors consistent with (c).

## Results

### UmbC proteins have paralogous interaction partners

To initiate our investigation of the UmbC proteins, we modeled their conserved domains using AlphaFold^17^. The ALF repeat portion of the proteins consistently adopted a ring structure, with interactions between ALF1 and ALF5 closing the ring and ALF4 and ALF8 located opposite (Figure 1b). The coiled-coiled domains of the proteins converge to form a stalk; in UmbC3 this stalk was predicted to extend unidirectionally the length of the domains, whereas the stalks of UmbC1 and UmbC2 adopted a bent configuration in initial models. Templating the models of UmbC1 and UmbC2 on UmbC3 using AlphaFold yielded straight stalks for these proteins, consistent with the modeled structures we obtained by AlphaFold of several other UmbC proteins (Figure S1b). Overall, the proteins adopt a lollipop-like structure approximately ∼300 Å in length.

The UmbC structure we predicted is dissimilar to characterized proteins and thus does not suggest how these proteins could function as polymorphic toxins. However, we reasoned that the ring arrangement of ALF repeats could serve as a platform for interaction with other proteins. To identify potential UmbC interaction partners, we generated *S. coelicolor* strains expressing C-terminally epitope-tagged UmbC1-3 from their native loci. Immunoprecipitation followed by mass spectrometry (IP-MS) revealed candidate interaction partners for each UmbC protein (Figure 1c, Table S3). Sequence comparison of the proteins established two families, which we named UmbA and UmbB. We noted that each UmbC is encoded proximal to a *umbA* gene and the gene encoding the UmbB proteins it precipitates (UmbA1-3, UmbB1-3) (Figure 1d). We also identified three UmbA proteins encoded outside of these regions (UmbA4-6); these proteins co-precipitated with each UmbC protein. The UmbC1 immunoprecipitation additionally yielded an Imm1 immunity protein family member, which we named UmbD1, as a candidate interaction partner. As observed for other polymorphic toxins, *umbD1* is located immediately downstream of its cognate toxin gene, *umbC1*. We did not identify candidate immunity proteins for UmbC2 or UmbC3 in our data; however, a gene encoding an Imm88 immunity family protein (UmbD3) is located downstream of *umbC3*.

### Protein interactions in the Umb complex

The UmbA proteins of *S. coelicolor* consist of a conserved N-terminal domain with high structural similarity to trypsin followed by a short helical linker to one (UmbA1-3, UmbA5, UmbA6) or more (UmbA4) sequence divergent domains predicted to function as lectins (Figure 2a, Figure S2a,b, and Tables S1,S4). With the exception of an intervening additional lectin domain in UmbA4, these domains belong to various β-propeller fold lectin families^18^. Unlike the UmbA proteins, UmbB proteins do not share significant sequence or predicted structural relatedness to characterized proteins. The predicted structure of these small proteins consists of an extended N-terminal disordered region linked by a short helix to a 10-stranded β-sandwich (Figure 2b).

**Figure 2:**
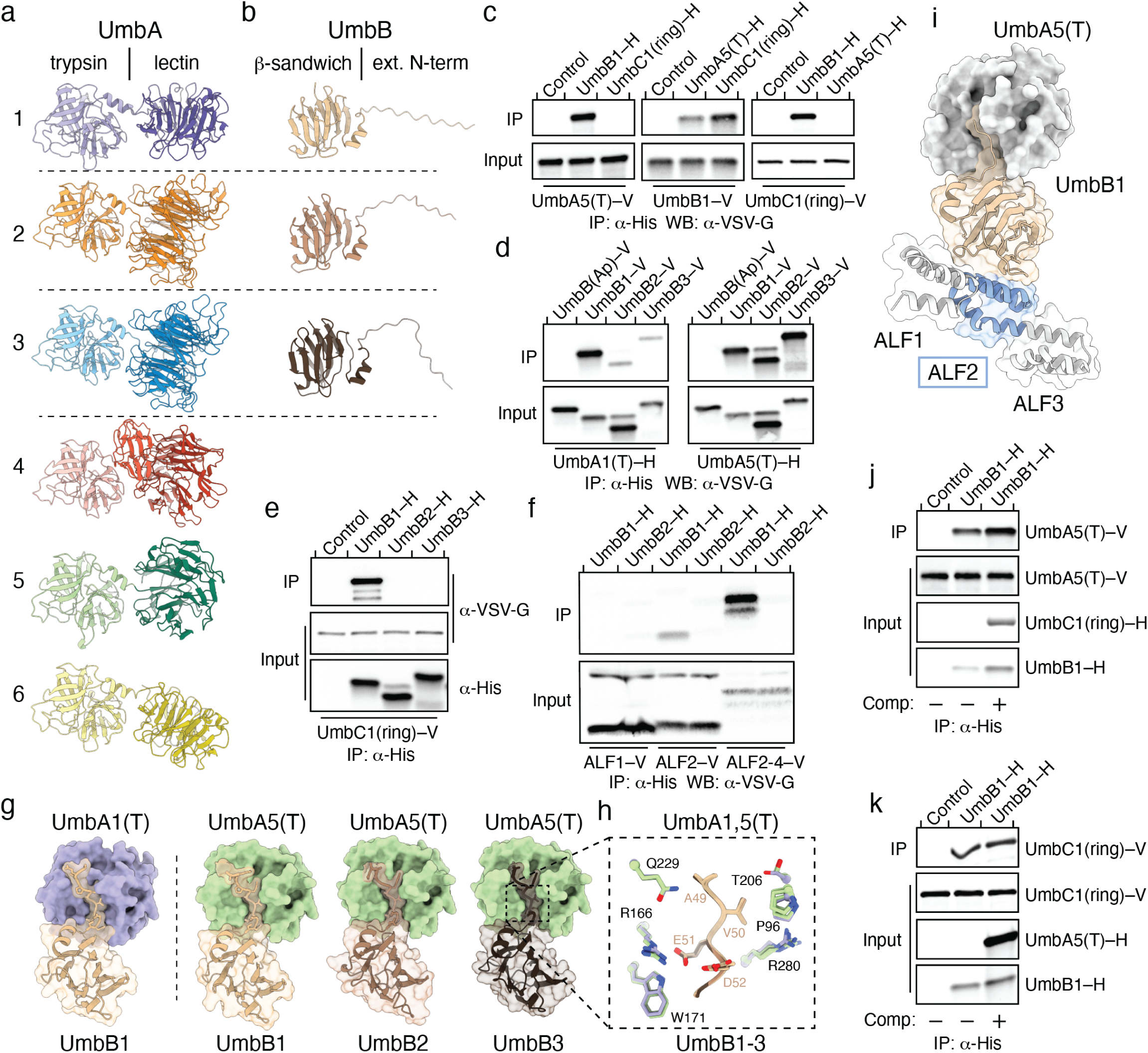
Protein-protein interactions in the Umb complex. **a,b** Predicted structural models for UmbA1-5 (a) and UmbB1-3 (b) of *S. coelicolor.* Dashed lines separate pairs of proximally-encoded proteins. **c-f,** Western blot (WB) analyses of immunoprecipitation (IP) experiments between the indicated heterologously expressed, tagged (–H, hexahistidine; –V, VSV-G epitope) Umb proteins. Controls lanes correspond to beads in the absence of a bait protein. UmbB(Ap) is a UmbB protein from the distantly related species *Actinoplanes philippinensis.* Additional input blots provided in Figure S3. **g,h,** Alphafold multimer-generated model for the interaction between the indicated UmbA and UmbB proteins of *S. coelicolor*, with surface representation highlighting the consistent predicted insertion of the N-terminus of UmbB proteins into the major cleft of UmbA trypsin domains. Additional predicted N-terminal disordered residues of UmbB1-3 are removed for clarity. Inset in (h) depicts strictly conserved residues in UmbA and UmbB in proximity to the modeled interaction interface. Side chains colored as in (g) and numbering corresponds to positions in UmbA5 and UmbB3. **i,** Ternary complex combining Alphafold multimer models of UmbB1–UmbA5(T) and UmbB1–ALF2 of UmbC1. Flanking ALF-repeats in UmbC1 (grey) are shown for context. **j,k**, Western blot analysis of competitive binding experiments between UmbB1 and its partners UmbA5(T) and UmbC1(ring). Purified competitor (Comp) UmbC1(ring)-H (j) or UmbA5(T)-H (k) were added in excess to IP experiments involving UmbB1 and UmbA5(T) or UmbC1(ring), respectively.

Next we sought to interrogate protein–protein interactions (PPIs) between predicted Umb complex components. In these experiments, we focused on the trypsin domains of UmbA proteins (UmbA(T)), given the likely involvement of their C-terminal lectin domains in carbohydrate binding and the challenges we encountered expressing their full-length form. Based on the assumption that PPIs involving UmbC would localize to the ALF repeats, we generated a DNA construct fusing the two sets of four repeats of UmbC1 into a ring, which removed the coiled-coil and C-terminal domains (UmbC1(ring)) (Figure S2c). Heterologous expression and co-immunoprecipitation studies provided evidence for direct interaction of UmbB1 with UmbA1(T), UmbA5(T), and UmbC1(ring) (Figure 2c,d and Figure S3). Consistent with our *S. coelicolor* UmbC immunoprecipitation findings, UmbA1(T) co-precipitated more robustly with UmbB1 than with UmbB2 or UmbB3, whereas UmbA5(T) co-precipitated similarly with UmbB1-3 (Figure 2d). Neither UmbA1(T) nor UmbA5(T) co-precipitated with an UmbB protein from the distantly-related organism *Actinoplanes philippinensis*.

In the UmbC ring, ALF repeats 1 and 5 are predicted to bind each other, apparently providing interactions important for uniting the two ring halves. Consequently, these repeats adopt an orientation and present a solvent accessible surface distinct from that of the other repeats (Figure S4). We reasoned that if ALF repeats mediate UmbB binding to the UmbC, this distinction would manifest as differential UmbB binding. Dissecting the UmbB–UmbC interaction, we found that UmbC1 displays specificity for UmbB1, and that ALF2, but not ALF1, is sufficient to mediate this interaction (Figure 2e,f). Furthermore, a construct composed of ALF2-4 co-precipitated more efficiently with UmbB1 than the single ALF2 repeat, suggesting that multiple ALF repeats engage UmbB (Figure 2f, Figure S3).

With experimentally determined PPIs between Umb proteins, we turned to AlphaFold to model their complexes. Strikingly, despite the sequence divergence between UmbB1-3 (39% average identity) and the trypsin domains of UmbA1 and UmbA5 (43% identity), the models consistently placed the extended N-terminal strands of UmbB1-3 into the prominent cleft of interacting UmbA proteins (Figure 2g). In this configuration, a consensus tetrapeptide motif within the UmbB proteins (Ala-Val-Glu-Asp) contacts conserved UmbA residues lining their prominent groove, which corresponds to the substrate binding cleft of trypsin proteins (Figure 2h). One particularly strong predicted contact is a salt bridge between the Glu within this motif and Arg156 or Arg166 of UmbA1 or UmbA5, respectively. Non-conservative substitutions at these positions in UmbB1 and UmbA5 abrogated their interaction (Figure S5a). Despite the small size of UmbB, modeling suggested the surfaces of UmbB1 that mediate UmbA and UmbC1 (ALF2) binding are non-overlapping (Figure 2i). This was supported by our finding that excess UmbA5 or UmbC1 did not interfere with UmbC1 or UmbA5 binding to UmbB1, respectively (Figure 2j,k).

Trypsin proteases utilize a Ser–His–Asp catalytic triad^19^. Alignment of UmbA1-6 with representative trypsin proteins showed that while the proteins share considerable sequence homology, no UmbA from *S. coelicolor* possesses the complete catalytic triad (Figure S5b). Moreover, we failed to detect catalytic activity from the purified trypsin domains of UmbA1 or UmbA5 using a universal trypsin substrate (Figure S5c,d). These data suggest that UmbA proteins utilize the trypsin fold in a non-canonical fashion, to bind, but not cleave, the extended N-terminus of their partner, UmbB. This mode of binding appears to permit promiscuity in UmbA–UmbB interactions and leave significant surface area of UmbB available for interaction with its other binding partner, UmbC.

### Structure of the Umb1 particle

The network of PPIs we uncovered between Umb proteins, combined with the multiplicity of ALF repeats in UmbC, suggested that the proteins could assemble into a large, multimeric particle. We found that relative to UmbC2 and UmbC3, UmbC1-based affinity purifications were homogenous and high yielding; however, instability near the C-terminal tagging site motivated us to identify the C-terminus of UmbA1 as an alternative site for isolating the complex by affinity chromatography (Figure 3a and Figure S6a). Isolation of UmbA1 from the supernatant of a *S. coelicolor* strain expressing a C-terminally octahistidine-tagged allele of the protein from its native chromosomal locus, followed by separation by size chromatography, yielded a complex composed predominantly of UmbA1, UmbA4-6, UmbB1 and UmbC1 (Figure S6b). Transmission electron microscopy (EM) of this negative-stained sample revealed that Umb1 particles adopt an umbrella-like morphology, leading to our naming these umbrella (Umb) toxin particles (Figure 3b and Figure S6c). The long, slender stalk of these particles extends ∼300 Å, whereas their crown has a width of ∼250 Å.

**Figure 3:**
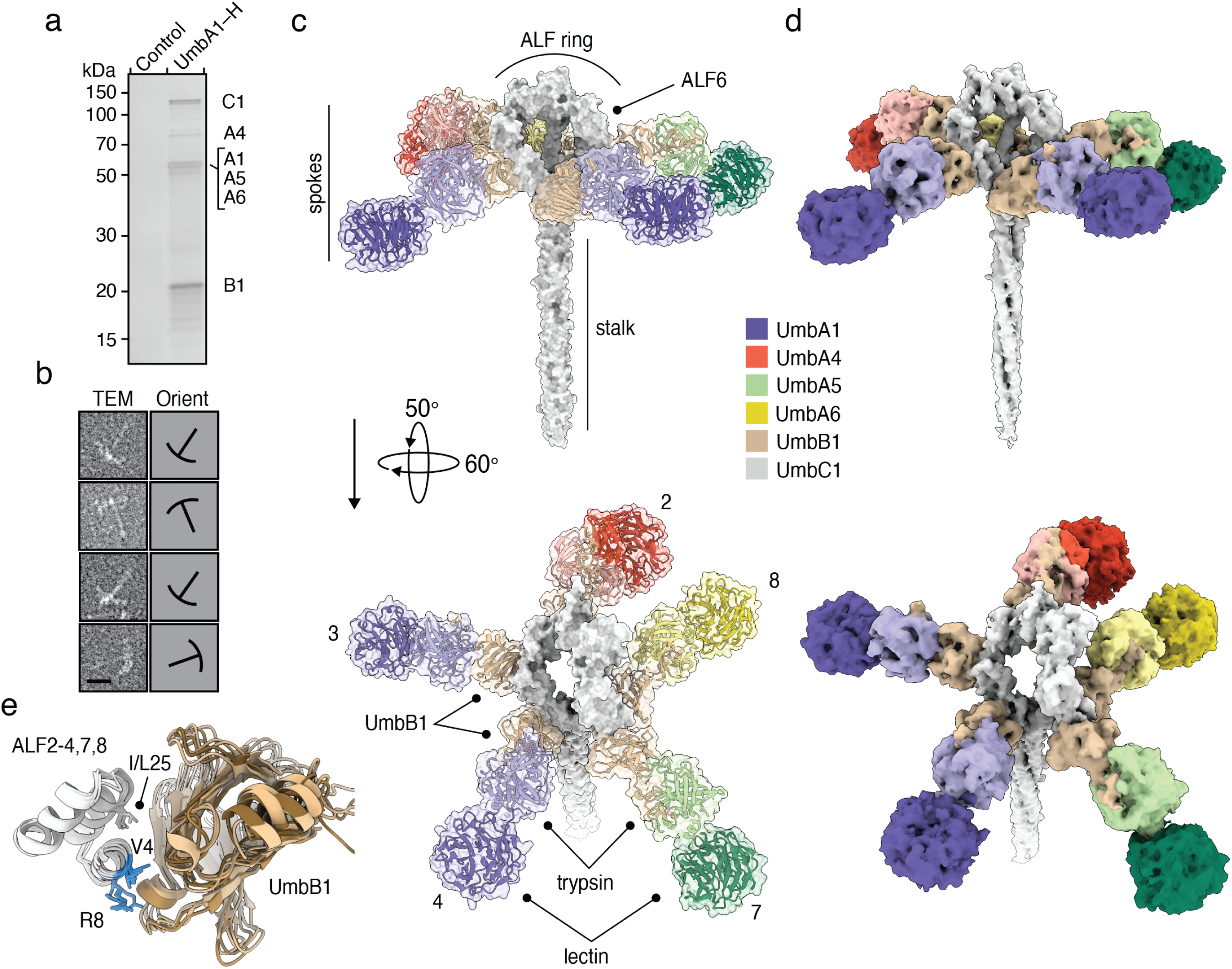
Structure of the Umb1 particle. **a,** Silver-stained SDS-PAGE analysis of the Umb1 protein complex purified from culture supernatant of *S. coelicolor.* **b**, Transmission electron microscopy (TEM) analysis of negative stained, purified Umb1 particles. Outlines indicating particle orientation (Orient) shown at right. Complete micrograph provided in Figure S6. **c,d,** Model (c) and 5.1 Å cryo-EM map (d) of the Umb1 particle. UmbA proteins modeled based on their relative abundance in the particle as measured by mass spectrometry. Spoke numbers correspond to the interacting ALF-repeat of UmbC1. ALF repeats 1, 5, and 6 (indicated) do not interact with UmbB1. The C-terminal domains of UmbC1 were not resolved in our structure. **e,** Superimposition of UmbB1-binding ALF repeats (white) in complex with their corresponding UmbB1 protomer (brown shades), extracted from the Umb1 particle structure. The predicted positions of the side chains of highly conserved residues in UmbB-interacting ALF repeats are shown for visualization purposes (but were not included in the final model as they are not resolved due to the limited resolution of the cryoEM map). Those not conserved in ALF6 are colored blue. Numbers reflect position within the repeat.

Using single particle cryo-EM, we obtained a structure of the Umb1 complex to a resolution of 5.1 Å using the gold-standard Fourier shell correlation cutoff of 0.143 (Figure S7a, Table S5). Our resolution was limited by particle heterogeneity and protein aggregation under cryogenic conditions (Figure S7b). The maps we obtained, in conjunction with Alphafold-predicted structures and our PPI analyses, allowed us to assemble a high-confidence model of the crown and proximal stalk regions of the Umb1 complex (Figure 3c,d and Figure S7c,d). The C-terminal toxin and HINT domains of UmbC1, which based on our model would localize to the tip of the stalk, were not resolved in our map. This could be the result of cleavage, mediated either by the HINT domain^20^ or an unknown protease, or may be due to flexible stretches of amino acids preceding these domains.

Our model revealed five spokes extending from the UmbC1 ring. Each spoke consists of an UmbB1–UmbA complex, connected to the ring via UmbB1 interaction with a single ALF repeat. Based on their relative abundance in our mass spectrometry data (Table S3), we modeled UmbA1 proteins in two spokes and UmbA4-6 in single spokes (Figure 3c,d). The lectin domains of the UmbA proteins reside at the distal ends of the spokes, a position compatible with engaging target cell receptor(s). The cryo-EM structure of the Umb1 complex confirmed that UmbC ALF repeats 1 and 5 do not bind UmbB1. Unexpectedly, it also revealed that ALF6 is not bound by UmbB1, producing a particle with five spokes rather than six (Figure 3c). We inspected the ALF–UmbB1 interface in order to identify the molecular basis of this selectivity. In spite of substantial variability in their sequences, the ALF repeats bind UmbB1 at a stereotyped location, with residues in two of its short helical segments providing many key contacts (Figure 3e). Several positions within this region that are identical across each UmbB1-binding ALF repeat and are predicted to mediate strong interactions with UmbB1, differ in ALF6 (Figure 3e). Most notably, repeat positions four and eight in ALF6 bear polar and acidic residues rather than the non-polar and basic residues, respectively, in the UmbB1-binding ALF repeats (Figure S8a). Based on our structure, we predict that these substitutions would preclude UmbB1 binding to ALF6. Indeed, high confidence AlphaFold models of UmbB1 in association with UmbB1-binding ALFs closely resemble those observed in our refined structure, whereas we were unable to obtain a high confidence model of UmbB1 bound to ALF6 or the other non-UmbB1 binding repeats (ALF1 and ALF5) using the program (Figure S8b). A similar trend was observed with the ALF repeats of UmbC2 and UmbC3 with UmbB2 and UmbB3, respectively, suggesting that the lack of UmbB binding to ALF6 may be a general feature of Umb particles. Our structure highlights UmbB1 as a remarkable adaptor protein and keystone component of the Umb toxin particle; it interacts with five sequence-divergent ALF repeats on one face and four different UmbA proteins on another. We are unaware of another characterized protein that displays this degree of binding partner plasticity.

### A Umb toxin potently and selectively targets *Streptomyces* spp

Functional predictions for the toxin domains associated with UmbC led us to speculate that umbrella particles act on bacterial targets. Indeed, heterologous expression of the C-terminal domains of the UmbC proteins of *S. coelicolor* led to a significant drop in bacterial viability (Figure 4a). The toxin domain of UmbC1 was particularly potent in these assays, and we confirmed the capacity of this predicted cytosine deaminase to introduce widespread C•G-to-T•A mutations in the DNA of intoxicated cells (Figure S9a-d). However, preliminary experiments measuring the impact of our purified Umb1 particle on the growth of a limited number of candidate bacteria did not identify clear targets of the toxin. To more broadly screen for Umb targets, we generated large quantities of concentrated Umb particle-enriched supernatant (Umb supernatant) from late exponential phase cultures of wild-type *S. coelicolor* and a control strain bearing deletions in each *umb* locus (Δ*umb* supernatant) (Figure S10a). This time point was chosen to maximize Umb particle levels based on prior genome-wide transcriptomic studies^21,22^. Next we used this material to screen for toxin targets among a collection of 140 diverse bacteria. Given the propensity of polymorphic toxins to act on closely related organisms, we included an abundance of *Streptomyces* spp. and other Actinobacterial species in our screen. This screen identified two candidate target species of the Umb toxin particles of *S. coelicolor* (Z score > 2.0), both of which are other *Streptomyces* species: *S. ambofaciens* (three strains) and *S. griseus* (Figure 4b, Figure S10b and Table S6). Subsequent time course experiments with these species and a control strain not hit in our screen demonstrated the capacity of *S. coelicolor* Umb supernatant to fully and specifically inhibit target cell growth in a manner dependent on Umb toxins. (Figure 4c and Figure S10c).

**Figure 4:**
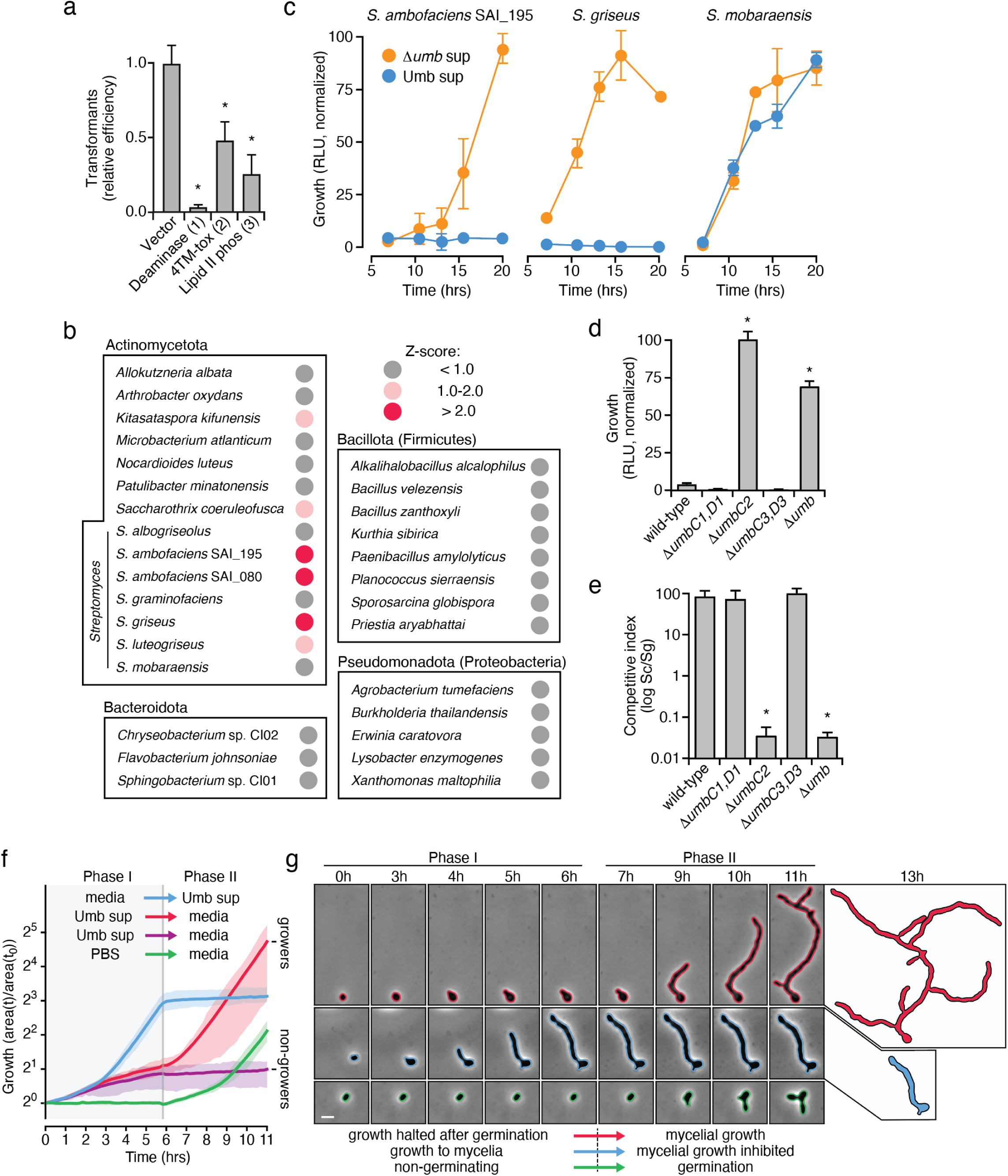
A Umb particle inhibits vegetative mycelial growth of specific *Streptomyces* species. **a,** Transformation efficiency in *Staphylococcus aureus* of plasmids expressing the indicated UmbC toxin domains relative to a vector control. The deaminase and Lipid II phosphatase (phos) domains were derived from UmbC1 and UmbC3 of *S. coelicolor.* The 4TM-toxin domain tested belongs to the same family as that from UmbC2 of *S. coelicolor*, but derives from *S. anulatus*, which encodes an adjacent immunity determinant necessary for generating the toxin expression construct. Means and standard deviations from at least two biological replicates with three technical replicates each are shown. Asterisks indicate transformation efficiencies significantly lower than the control (p<0.01, Dunnett’s multiple comparison test). **b**, Select Umb toxin target screening results. Z-scores calculated from relative growth (as determined by ATP quantification) in the presence or absence of Umb toxins; scores >2 indicate significant Umb-dependent inhibition. Additional strains screened are shown in Data Figure S10b and raw data are provided in Table S6. **c,** Growth of the indicated target and non-target strains treated with Umb or Δ*umb* supernatant (10% (v/v)) as determined by luminescence-based ATP quantification (RLU, relative luminscence units). Colony forming units were quantified at the 16 hr time point (Figure S10c). **d,** Growth yields of *S. griseus* treated with *S. coelicolor* Umb supernatant from the indicated *S. coelicolor* strains. Growth was measured after 16 hr as in (c). **e**, Outcome of growth competition assays between the indicated strains of *S. coelicolor* and *S. griseus.* Means and standard deviations from technical triplicates are shown in c-e, and asterisks (d,e) indicate differences significant from wild-type controls (p<0.01, Dunnett’s multiple comparisons test). **f, g,** Single cell-based microscopic analysis of aggregate (f) or representative (g) *S. griseus* growth as determined by cell area during exposure to the indicated treatments in a microfluidic flow cell. Cells receiving Umb supernatant during the phase I fell into two classes: cells able to resume growth upon the infusion of toxin-free medium (growers) and those remaining arrested (non-growers). Data for individual cells from these and other treatment groups provided in Figure S11 and in Video S1. Shading indicates interquartile ranges. Red, n = 41; Blue, n = 55; Purple, n = 29; Green, n = 62. **g**, Cropped micrograph regions showing representative cells, outlined with Omnipose-generated segmentation masks, from the indicated treatment groups in (f). At 13 hr, only cell masks are presented. Scale bar, 2 μM.

The *S. griseus* strain hit in our screen is a type strain that is amenable to genetic manipulation and straightforward to cultivate^23^. We thus selected this target organism to further characterize Umb-dependent toxicity. To identify the Umb particle(s) responsible for inhibiting *S. griseus*, we tested the toxicity of Umb supernatant deriving from *S. coelicolor* strains unable to synthesize individual Umb particles. Inactivation of *umbC2*, but not *umbC1* or *umbC3*, abrogated Umb supernatant growth inhibitory activity toward the organism (Figure 4d). We next performed growth competition experiments to determine whether the level of Umb2 produced by *S. coelicolor* during co-culture is sufficient to intoxicate target cells. Strikingly, we found that an *S. coelicolor* strain lacking Umb2 function is >1,000-fold less fit than the wild-type in co-culture with *S. griseus* (Figure 4e). In summary, these data show that the secreted Umb toxins of *S. coelicolor* potently inhibit the growth of other Streptomyces species.

### The Umb2 particle inhibits mycelial growth

Streptomyces undergo a complex developmental program, proceeding from spore germination to vegetative mycelial growth, followed by production of aerial mycelia and sporulation. To gain insight into the possible ecological role of Umb toxin particles during competition between *Streptomyces*, we sought to determine the developmental stage at which target *Streptomyces* species are susceptible to Umb particle-mediated intoxication. Single cell level analysis of time-lapse microscopy data revealed that Umb supernatant from wild-type *S. coelicolor* does not impact spore germination in the Umb2 target *S. griseus* (Figure 4f,g, Figure S11, and Video S1). Rather, like spores treated with media or Δ*umb* supernantant, those treated with Umb supernatant increase in size and elaborate nascent germ tubes – phenomena not observed under conditions non-permissive to germination. However, spores treated with media or Δ*umb* supernatant completed germination and formed mycelia, while Umb supernatant-treated cells arrested at the nascent germ tube phase (Figure 4f,g and Video S1). Upon replacement of the Umb supernatant with media, a proportion of the population resumed vegetative growth after a variable lag period, while other cells remained inhibited (Figure S11). We speculate that the vegetative bacterial surface area exposed to the Umb particle during germination determines the dose of toxin received, and thus influences the subsequent fate of the cell.

Our data also revealed that the addition of Umb supernatant to actively growing mycelia produces an immediate, complete and persistent growth arrest (Figure 4f,g and Video S1). We did not observe lysis of intoxicated cells, consistent with the predicted pore forming activity of UmbC2. Together, these results demonstrate that the Umb2 particle acts specifically to inhibit vegetative mycelial growth of target organisms. Transcriptomic studies and our proteomics data show that Umb toxins are also produced during this phase of the *Streptomyces* lifecycle, suggesting a physiological function in mediating the outcome of competition between populations of vegetatively growing *Streptomyces*^21,22^. This is distinct from small molecule antimicrobials produced by Streptomyces, which generally target a much broader group of organisms for the purpose of limiting access to nutrients released by lysed kin cells during aerial hyphae formation^5^.

### Diversity and distribution of Umb toxins

The Umb particles of *S. coelicolor* confer a significant advantage in competition with multiple species. Given the prevalence of antagonistic interactions among bacterial species, we reasoned that others might harbor and utilize Umb toxins in an analogous fashion. Leveraging our *S. coelicolor* findings pertaining to the particle constituents and genetic organization of Umb1-3, we searched publicly available bacterial genomes to broadly define the distribution of Umb toxins. In total, we identified 1,117 genomes, deriving from 875 species, that we predict possess the capacity to synthesize one or more Umb particles (UmbB and UmbC within 10 genes of each other) (Table S1). Over half of these correspond to species within the order Streptomycetaceae; the remaining *umb* loci-containing species distribute among six other orders of Actinobacteria (Figure 5a). In multiple bacteria capable of synthesizing distinct Umb particles, we identified UmbA proteins encoded at loci unlinked to those encoding UmbB and UmbC (Figure 5b). This suggests that the association of “orphan” UmbA proteins with multiple particles, as observed in *S. coelicolor*, may be commonplace. It is notable that we did not find support for Umb particle production by bacteria outside of Actinobacteria. If the action of Umb toxins is restricted to related species or to bacteria that exhibit mycelial growth, this finding could reflect the phylogenetic limits of targeting by this mechanism.

**Figure 5:**
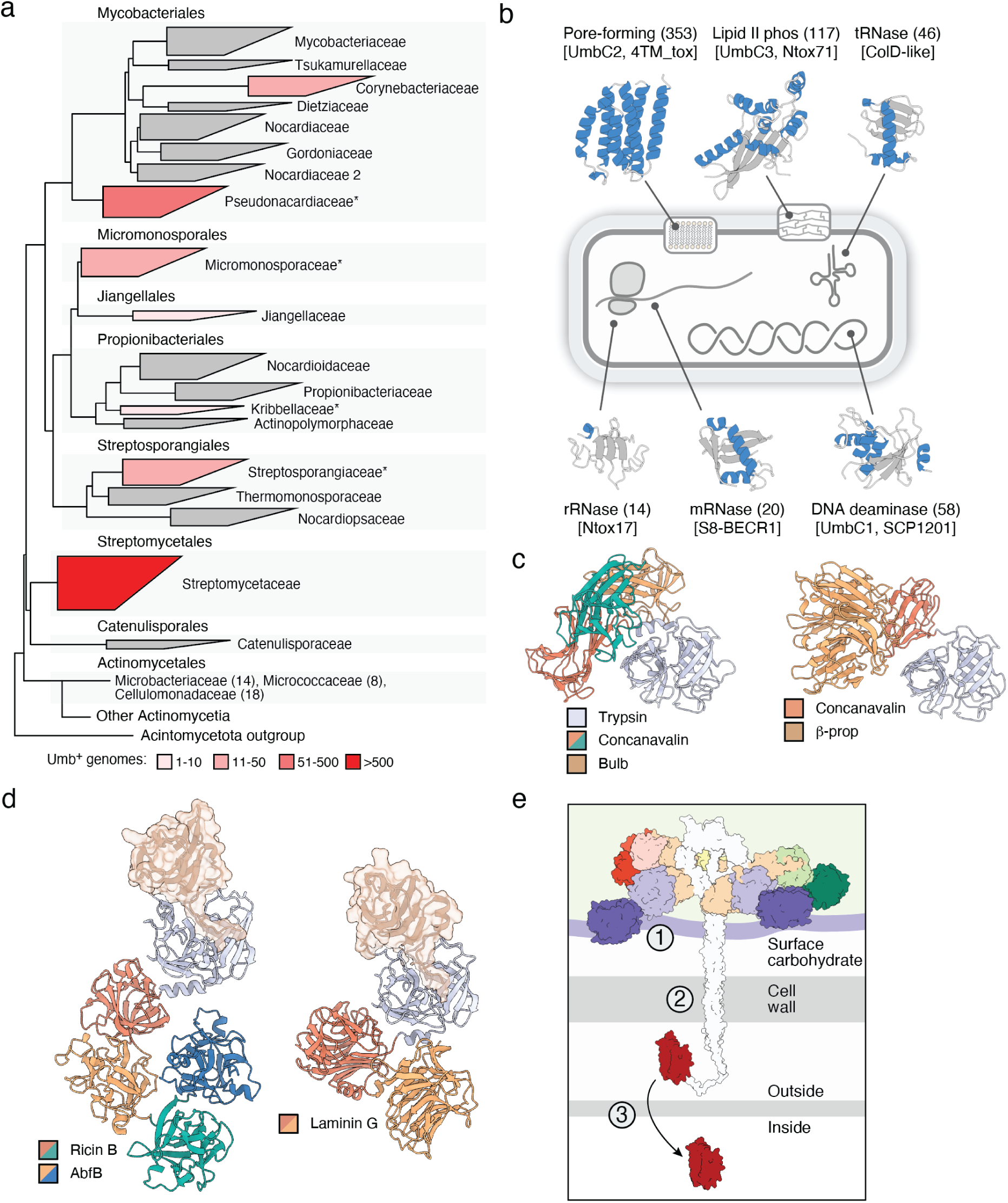
Phylogenetic distribution and functional diversity of Umb proteins. **a,** Phylogenetic tree of orders and families within Actinomycetia, colored to indicate the number of genomes positive for Umb toxin particle loci. Within Actinomycetales, only those families containing *umb* loci are listed, with the number of *umb*-containing genomes in parentheses. Asterisks indicate families for which representative *umb* loci are shown in Figure S12. **b,** Schematic indicating the molecular targets of select toxin domains commonly found in UmbC proteins and representative models for the domains generated with AlphaFold. Models colored by secondary structure (blue, α-helices; grey, loops and β-strands). Values in parentheses indicate the number of UmbC proteins we detected carrying the indicated toxin domain. Toxin family names are provided in brackets and in Table S2. **c,d,** Predicted structural models of example UmbA proteins selected by virtue of containing multiple distinct or repeated lectin domains (c) or fusions between UmbA and UmbB proteins (d). The UmbB domains of bifunctional UmbAB proteins in (d) are shown in transparent surface representation and in the same orientation to highlight their conserved interaction with the major cleft of the trypsin-like domain. β-prop, UAL-Bprop-1 family identified in this study, see Table S4. (e) Model for the intoxication of target cells by Umb toxins, highlighting outstanding questions. These include the identity of receptor(s) on target cells and the involvement of the lectin domains in mediating binding (1), the role of the stalk in toxin delivery (2), and the mechanism of toxin translocation into target cells (3).

We found 77 divergent toxin families associated with the UmbC proteins identified in our analyses (Table S2). While many of these bear sequence similarity to toxin domains associated with other polymorphic toxin systems, many, including the two most frequently observed in UmbC proteins, represent previously unrecognized families (4TM_tox, Ntox71). Functional predictions suggest that as a group, Umb toxins act upon a striking range of essential cellular processes (Figure 5c and Table S2).

A unique feature of Umb particles uncovered by our work in *S. coelicolor* is their incorporation of variable lectin domains via promiscuous UmbA binding. Taken together with their accessibility at the ends of Umb particle spokes, we hypothesize that these domains mediate target cell binding and, at least in part, underpin the species intoxication selectivity we observe. Examination of the 882 UmbA proteins identified by our search highlighted extraordinary family- and within family-level diversity in the lectin domains associated with these proteins (Table S4). Moreover, we identified striking structural diversity among UmbA proteins, including those that, like *S. coelicolor* UmbA4, encode multiple distinct lectin domains, and others that are fused to UmbB-like domains (Figure 5d,e). AlphaFold models of the latter predict that, despite their fusion, the predominant engagement mode of the two domains mirrors that which we identified for the individually encoded proteins; an extended N-terminal structure of the UmbB domain inserts within the major cleft of the trypsin-like domain. Taken together, the diversification of toxin and lectin domains associated with Umb toxin particles provides evidence for a molecular arms race between between producer and target cells, wherein target cells can escape intoxication either by receptor modification or by acquiring a downstream, direct toxin resistance mechanism.

## Discussion

Umbrella toxin particles represent a previously unrecognized component of the antibacterial arsenal of *Streptomyces*. We hypothesize that Umb particles mediate dynamic short-range antagonism between the vegetative mycelia of competing species vying for the same niche. This would provide the evolutionary pressure driving Umb particle selectivity and diversification, as the overlap in niches of highly related bacteria increases their probability of repeated encounters^2,7,24^. The chemical and biophysical properties of Umb particles are also consistent with this role. Umb toxin particle complexity and apparent vulnerability to proteases or other insults suggests they are short-lived, and thus unable to act at longer length scales. Indeed, these properties of the Umb particles may underlie why such potent toxins escaped detection for the more than 100 years that scientists have been studying antagonistic interactions between *Streptomyces* species^25^.

Polymorphic toxins are found in a wide range of organisms, function in many contexts, and access their targets through a diverse set of delivery systems^10,15^. Yet, it is difficult to identify a characterized polymorphic toxin system that represents a close analog of the Umb particle. In certain respects, colicins – antibacterial proteins produced by *E. coli* – might be considered most comparable. Like Umb particles, these are secreted proteins that mediate interactions among closely related strains^26^. Furthermore, seemingly analogous to Umb particles, they do so via assorted receptor binding domains coupled to polymorphic toxin domains^27^. However, there exists a multitude of features distinguishing colicins and Umb toxin particles, and even their few similarities are superficial. For example, colicins typically target strains belonging to the species of the producer cell, and the diversity of receptor protein binding domains in colicins (<10) is far eclipsed by the diversity of carbohydrate-binding lectin domains associated with Umb particles. Perhaps the starkest of differences between the two polymorphic toxins is their mechanism of secretion, which further highlights their apparently disparate physiological functions. Colicins access the extracellular milieu through a non-canonical mechanism that requires the action of bacteriocin release proteins, referred to as lysis or killing proteins for the death they inflict upon producer cells^28^. Colicin expression is thereby under the control of a repressor responsive to cellular damage and the utilization of these toxins can be categorized as an altrustic behavior^29^. On the other hand, UmbA-C each possess N-terminal Sec (UmbA,B) or TAT (UmbC) secretion signals and we find no data suggesting that the release of Umb particles is detrimental to producer cells. Moreover, our results and prior genome-wide transcriptome studies suggest that, at least in *S. coelicolor*, Umb particles are produced during log phase vegetative mycelial growth, well before the onset of widespread cell death^21,22^. Our work suggests that continued exploration of proteins containing polymorphic toxin domains in diverse bacteria may reveal additional structurally and mechanistically unprecedented toxins.

This work identified the Umb toxin components of *S. coelicolor*, defined their pairwise interactions, revealed the ultra-structure of the structure of the particle they form, and it established the role of these particles in interbacterial antagonism between *Streptomyces* species. Nevertheless, important open questions for future studies remain. With regard to target cells, do the UmbA lectin domains play a role in recognition, what is the identity of the receptor(s), what role does the stalk play, and how do toxins with cytoplasmic targets cross the membrane (Figure 5e)? In the producer cell, key open questions include how are the *umb* genes regulated, how and where do Umb particles assemble, and are Umb particles from across Actinobacteria mounted universally to mediate interbacterial antagonism? It is also of interest to consider the potential biotechnological and therapeutic applications of Umb particles. *Mycobacterium tuberculosis* and *Corynebacterium diptheria* are two important human pathogens that, as Actinobacteria, are potential Umb targets, and for which resistance to traditional antibiotics is of growing concern^30,31^. In total, our work identifies an antibacterial toxin particle with promise to expand our knowledge of the mechanisms, ecological implications, and biotechnological applications of interbacterial antagonism.

## Methods

### Bacterial strains and culture conditions

A complete list of strains used in this study can be found in Table S7. *Escherichia coli* strain DH5α was used for plasmid maintenance, strain ET12567 (pUZ8002) for interspecies conjugation, and strain BL21 for protein expression. *E. coli* strains were grown in Lysogeny Broth (LB) at 37℃ with shaking or on LB medium solidified with 1.5% w/v agar. *Staphylococcus aureus* strain RN4220 was used for plasmid maintenance and protein expression. *S. aureus* was grown in B2 broth, LB supplemented with 0.2% (w/v) glucose (LBG), or on LBG solidified with 1.5% (w/v) agar. Strain *Streptomyces coelicolor* A3(2) was employed in Umb characterization studies. Unless otherwise noted, this and other *Streptomyces* species employed were cultivated in R5 or TSBY liquid medium at 28℃ in baffled flasks with glass beads (3mm diameter) shaking at 220 r.p.m. or on TSB, ISP2, ISP4, or SFM solidified with 1.5% w/v agar. Growth conditions of diverse bacterial species used in the broad Umb sensitivity screen can be found in Table S6. Media were supplemented as needed with antibiotics at the following concentrations: carbenicillin (150 µg ml^−1^, *E. coli*), apramycin (50 µg ml^−1^, *E. coli* and *Streptomyces*), kanamycin (50 µg ml^−1^, *E. coli*), gentamicin (15 µg ml^−1^, *E. coli*), trimethoprim (50 µg ml^−1^, *E. coli* and *Streptomyces*), chloramphenicol (25 µg ml^−1^, *E. coli*; 10 µg ml^−1^, *S. aureus*), and hygromycin (25 µg ml^−1^, *E. coli*).

### Plasmid construction

Plasmids used in this study, details of plasmid construction, and primers employed in this work are provided in Table S7. Primers and synthetic DNA fragments were obtained from IDT. All plasmid constructs were constructed using Gibson assembly, and all constructs were confirmed by sequencing. For heterologous expression of Umb complex proteins in *E. coli*, the genes were amplified and inserted into NcoI- and XhoI-digested pET-22b(+) or NdeI- and XhoI-digested pET-28b(+) to generate C-terminal or N-terminal hexahistidine fusions, respectively. VSV-G fusions, point mutations, and linkers were introduced to genes amplified from the *S. coelicolor* genome through the cloning primers. *umbC1(ring)* expression plasmids were constructed by amplifying *ALF1-4* (residues A14-A209) and *ALF5-8* (residues A500-H766) as two DNA fragments with a linker of two GGGGS repeats introduced in the cloning primers.

Plasmids used for heterologous expression of UmbC1 and UmbD1 in *E. coli* for mutational profiling were pSCrhaB2 and pPSV39-CV, respectively. To generate these plasmids, the genes were amplified from synthesized DNA fragments codon optimized for expression in *E. coli.* Plasmid pEPSA5 was used for heterologous expression of various *umbC* toxin domains in *S. aureus*. The toxin domain was either inserted into digested plasmid alongside an N-terminal 3xFLAG tag second insert fragment or alongside a signal sequence-containing second insert fragment, with an N-terminal 3xFLAG tag being introduced via the cloning primers. These Gibson reactions were transformed into *S. aureus* RN4220 via electroporation, and transformants were maintained in LB supplemented with 0.2% w/v glucose (to repress toxin expression) and chloramphenicol.

*S. coelicolor* genetic manipulation was conducted using a derivative of the suicide vector pKGLP2^32^, in which the hygromycin resistance cassette (*hyg*) was replaced with the apramycin resistance gene (*aac*(3)) and promoter from pSET152^33^. This plasmid, pKGLP2a, was generated by amplifying the vector backbone of pKGLP2 and the apramycin resistance cassette from pSET152 by PCR and combining by Gibson assembly. Constructs for introducing deletions, epitope tags and point mutations in the *S. coelicolor* genome with pKGLP2a were generated using Gibson assembly of 1.5-2 kb arms flanking the site of modification.

### Structural modeling of Umb proteins and protein-protein interactions

Structural predictions for UmbC1-3 were made using AlphaFold2^17^. MSAs were generated by running hhblits^34^ against UniRef30^35^ and BFD^36^. These MSAs were uploaded to ColabFold^37^ and a total of five AlphaFold predictions were generated for each target. Only UmbC3 generated predictions that were consistent with the cryoEM density of the protein while models for UmbC1 and UmbC2 all resulted in the long coiled-coil folding back on itself. This prompted the decision to use the UmbC3 model as a template structure for predicting UmbC1 and UmbC2, which allowed the generation of models with a straight coiled-coil consistent with the cryoEM density. The models with highest predicted lDDT were selected for each.

RoseTTAFold2^38^ was used to predict UmbA:UmbB protein structures. MSAs were generated as described previously for UmbC1-3. Paired MSAs for all UmbA:UmbB pairs were generated by matching taxonomy IDs, following the published methods^39^. These paired MSAs were provided as inputs to RoseTTAFold2, and produced confident predictions in all cases (plDDTs>0.8). A similar method was used to compute predictions for interactions between UmbB and individual ALF repeats of UmbC1-3. In brief, MSAs were generated for UmbB1, UmbB2, UmbB3, UmbC1, UmbC2, and UmbC3 by running hhblits against Uniref30 and BFD, and paired MSAs for all three pairs were generated by maxing taxonomy IDs. Then, predictions were made for each UmbB model against each of the eight ALF repeats of the corresponding UmbC model. Rather than regenerating the MSA for individual repeats, the paired full-length MSA was trimmed over the region of each repeat.

Owing to the availability of cryoEM data, models for UmbC1:UmbB1 were generated first. Three different variants of repeat modeling were tried: a) trimming to exactly the two-helix repeat; b) extending by 5 residues on either side of the repeat; and c) extending by 10 residues on either side of the repeat. To evaluate each modeling variant, the predicted structure and predicted interface error (pAE) of the UmbC:UmbB interface^17^ were considered. All three trimming approaches yielded results consistent with the EM data, but the most distinct signal in terms of interfacial pAE was achieved by adding in 10 residues of padding. This strategy was applied to UmbC2:UmbB2 and UmbC3:UmbB3.

### Construction of genetically modified *S. coelicolor* strains

Genetic modification constructs in the pKGLP2a suicide plasmid were transferred to *S. coelicolor* by intergeneric *E. coli-Streptomyces* conjugation using donor strain *E. coli* ET12567 (pUZ8002) as described previously^40^. Briefly, overnight cultures of *E. coli* ET12567 (pUZ8002) harboring the plasmid to be transferred were grown in LB supplemented with chloramphenicol, kanamycin and apramycin. These cultures were washed, concentrated, and combined with *Streptomyces* spores following a 10-minute 50℃ heat shock treatment. The mixture was plated on SFM media supplemented with 10mM MgCl_2_ and incubated at 30℃ for 16-20 hours. The plate was then overlaid with 1 mL sterilized dH_2_O supplemented with trimethoprim and apramycin. Incubation was continued at 30℃ until transconjugants appeared and were re-streaked to media supplemented with trimethoprim and apramycin. Apramycin-resistant *S. griseus* was generated by intergeneric transfer of pSET152_aac(3)IV-bla using donor strain ET12567 (pUZ8002) by the same method.

### Immunoprecipitation and mass spectrometry analysis of UmbC-interacting proteins from *S. coelicolor*

Spores of *S. coelicolor* strains containing *umbC1*-VSV-G, *umbC3*-VSV-G or *umbA1*-VSV-G at the native loci were inoculated in R5 medium and grown for 36 h, then back diluted 1:200 in 50 mL R5 medium and further grown for 24-30 h until OD_600_ reached 3-4. Spores of *S. coelicolor* containing *umbC2*-VSV-G at the native locus were inoculated in 50 mL TSBY medium and grown for 36 h. For each strain, 10 mL of the cell culture was then mixed with 2.5 mL 5x lysis buffer (750 mM NaCl, 100 mM Tris-HCl pH 7.5, 10% glycerol [*v/v*], 1 mg mL^-1^ lysosome, and 1 mU benzonase). Cells were lysed by sonication and cellular debris removed by centrifugation at 35,000 x g. for 30 minutes. VSV-G tagged proteins were enriched by incubation of cell lysates with 40 μL of anti-VSV-G agarose beads at 4℃ for 4-5 h with constant rotation. The agarose beads were then pelleted by centrifugation at 300 x g for 2 minutes, washed three times with 10 mL wash buffer (150 mM NaCl, 2% glycerol, and 20 mM Tris-HCl pH 7.5), and then washed three times with 10 mL 20 mM ammonium bicarbonate. VSV-G agarose beads and bound proteins were then treated with 10 μL of 10 μg/μL sequence grade trypsin (Promega) for 16 h at 37℃ with mild shaking. After digestion, the agarose beads and peptides were mixed lightly and centrifuged at 300 x g for 2 min. After collection of the supernatant, 90 μL of 20 mM ammonium bicarbonate was added to the beads, mixed lightly and centrifuged again. The supernatant was collected and combined as the peptide fraction. The mixture was reduced with 5 mM Tris(2-carboxyethyl) phosphine hydrochloride for 1 h at 37℃, followed by alkylation using 14 mM iodoacetamide for 30 min in the dark at room temperature. The alkylation reaction was quenched by adding 5 mM 1,4-dithiothreitol. Acetonitrile (ACN) and trifluoroacetic acid (TFA) were added to the samples for a final concentration of 5% (v/v) and 0.5% (w/v), respectively. Then, the samples were applied to MacroSpin C18 columns (7-70 μg capacity) that had been charged with 100% ACN, LC-MS grade water and 0.1% TFA. Bound peptides were washed twice with 0.1% TFA and then eluted with 80% ACN with 25 mM formic acid (FA). The dried peptides were dissolved in 5% ACN with 0.1% FA and analyzed by LC-MS/MS as described previously^41^. Data were analyzed using MaxQuant^42^, and filtered to remove noise from low abundance proteins with five or fewer spectral counts in IP samples. Enrichment of proteins in the IP samples was determined by dividing the relative abundance of each protein passing the filtering criteria in the IP samples by its relative abundance in the control.

### Purification of heterologously-expressed Umb proteins

A subset of the protein-protein interaction studies and the protease activity assay we performed employed purified, heterologously expressed Umb proteins. To purify these proteins, overnight cultures of *E. coli* BL21 Rosetta 2 DE3 carrying pET-22b(+) or pET-28b(+) constructs expressing the protein of interest were back diluted 1:300 in 2xYT broth and grown at 37℃ shaking at 220 r.p.m. until OD_600_ = 0.4. The incubation temperature was lowered to 18℃; after 30 minutes, IPTG was added to a final concentration of 0.3 mM and the cultures were incubated for a total of 18 hours. Cells were then collected by centrifugation and resuspended in lysis buffer containing 200 mM NaCl, 50 mM Tris-HCl pH 7.5, 10% glycerol (v/v), 5 mM imidazole, 0.5 mg/mL lysosome, and 1 mU benzonase. Cells were then lysed by sonication and cellular debris removed by centrifugation at 35,000 x g for 30 minutes at 4℃. The 6xHis-tagged proteins were purified from lysates using a 1 mL HisTrap HP column on an AKTA fast protein liquid chromatographer (FPLC). Column-bound protein was eluted using a linear imidazole gradient from 5 mM to 500 mM. Protein purity was assessed by SDS-PAGE and Coomassie staining. The fractions with high purity were concentrated using 10 kDa cutoff Amicon filters and then further purified by FPLC using a HiLoad^TM^ 16/600 Superdex^TM^ 200 pg column (GE Healthcare) equilibrated with sizing buffer (500 mM NaCl, 50 mM Tris-HCl pH 7.5, 10% glycerol [v/v]).

### Protein-protein interaction assays

Interactions between Umb proteins were probed using proteins heterologously expressed in *E. coli*. For tests of the interactions between UmbB1, UmbA5(T), and UmbC1(ring), 400 μL equilibration buffer (200 mM NaCl, 50 mM Tris-HCl pH 7.5, 10 mM imidazole) containing with 5 µg of purified UmbB1-H, UmbA5(T)-H, or UmbC1(ring)-H was mixed with 400 μL *E. coli* cell lysate containing UmbA5(T)-V, UmbC1(ring)-V, or UmbB1-V, respectively. To assess input protein levels, 40 μL of these samples were mixed with 4x Laemmli loading buffer (Bio-Rad) and boiled 20 min at 95 ℃ for Western blot analysis. The remaining protein mixtures were incubated with 50 μL Ni-NTA agarose beads (QIAGEN) at 4℃ for 1.5 h with constant rotation. Agarose beads were pelleted by centrifugation at 300 × g for 3 min and washed five times with 1.4 mL wash buffer (500 mM NaCl, 50 mM Tris-HCl pH 7.5, and 25 mM imidazole). Proteins bound to the Ni-NTA resin were then eluted by 100 μL elution buffer (500 mM NaCl, 50 mM Tris-HCl, and 300 mM imidazole). The eluate was mixed with 4 × Laemmli loading buffer, boiled and subjected to Western blot analysis. For the other protein-protein interaction assays, *E. coli* cell lysates containing 6xHis-tagged bait proteins were mixed directly with *E. coli* cell lysates containing VSV-G tagged target proteins, then incubated with Ni-NTA agarose beads, washed and processed as above. For the competitive binding experiments between UmbB1 and its partners UmbA5(T) and UmbC1(ring), 3 μg of purified UmbB1-H was incubated with 50 μL Ni-NTA agarose beads at 4℃ for 1 h with constant rotation, followed by two washes with equilibration buffer. 400 μL equilibration buffer with 2-fold molar excess of purified competitor UmbC1(ring)-H or UmbA5(T)-H was mixed with 400 μL *E. coli* cell lysates containing UmbA5(T)-V or UmbC1(ring)-V, respectively. The protein mixture was further incubated with UmbB1-H bound to Ni-NTA agarose beads, and then washed and processed as above.

### Western blot analysis

To analyze the protein-protein interaction assays performed with heterologously expressed Umb proteins, equal volumes of input samples or Co-IP samples were resolved using SDS-PAGE, then transferred to nitrocellulose membranes. Following the transfer, membranes were blocked in in TBST (10 mM Tris-HCl pH 7.5, 150 mM NaCl, and 0.1% w/v Tween-20) with 5% w/v bovine serum albumin (BSA) (RPI CAS #9084-46-8) at room temperature for 1 hr. Primary antibodies (α-His HRP conjugated (Qiagen 34460) or α -VSV-G (Millipore sigma V4888-200UG)) were then added at a dilution of 1:5000 and incubated at room temperature for 1 hour. Blots were then washed four times with TBST, and anti-VSV-G blots were incubated with secondary antibody (α -Rabbit HRP conjugated (Sigma Aldrich, A6154-1ML)) diluted 1:5000 in TBST at room temperature for 1 hr. Finally, blots were washed four times with TBST again and were developed using Clarity Max Western ECL Substrate (Bio-Rad Cat # 1705062) and visualized using the Invitrogen iBright 1500 imager.

### Trypsin assays

The protease activity of purified UmbA1 and UmbA5 trypsin domains was assessed using Roche’s universal protease substrate following the manufacturer’s protocol. Briefly, 50 μL substrate solution (0.4% casein), and 50 μL incubation buffer (0.2 M Tris-HCl pH 7.8, 0.02 M CaCl_2_) were combined with 100 μL sample buffer (300 mM NaCl, 50 mM Tris-HCl pH 7.8) containing either 500 ng purified protein (UmbA1(T) or UmbA5(T)), 100 ng trypsin (positive control), or no protein (blank). The mixture was incubated at 37℃ for 15 minutes before adding 480 μL stop reagent (5% Trichloroacetic acid [w/v]).. The samples were further incubated 37℃ for 10 minutes and centrifuged at 13,000 x g for 5 minutes. 400 μL of the reaction mixture was then combined with 600 μL assay buffer (0.5 M Tris-HCl, pH 8.8) in a cuvette and absorbance was measured at 574 nm.

### Purification of the Umb1 particle for structural studies

*S. coelicolor* spores expressing UmbA1-8xHis from the native locus were inoculated into 30 mL R5 media and incubated at 30℃ shaking at 220 r.p.m. for 36 hours. Cultures were back diluted 1:200 in 50 mL R5 for a total combined culture volume of 700 mL and incubated 24-30 hours, until OD_600_ reached ∼4.. Cells were then pelleted by spinning at 21,000 x g for 45 minutes and the resulting supernatant was filtered (GenClone 25-229, Vacuum Filter Systems, 1000ml PES Membrane, 0.22µm). Next, 600 mL supernatant was combined with 150 mL 5x lysis buffer (1 M NaCl and 250 mM Tris-HCl pH 7.5) and run over a 1 mL HisTrap FF column on an AKTA FPLC purification system to purify the His-tagged proteins. The bound proteins were eluted using a linear imidazole gradient from 0 mM to 300 mM. Collected fractions were pooled and concentrated using a 100 kDa cutoff Amicon concentrator until reaching a final volume of ∼600 μL. The protein sample was further purified by FPLC using a Superose^®^ 6 Increase 10/300 GL column (GE Healthcare) equilibrated in sizing buffer (150 mM NaCl, 20 mM Tris-HCl pH 7.5, and 3% glycerol). Each fraction was assessed for purity by SDS-PAGE and silver staining. The fractions with the highest purity and concentration were used for negative stain EM or CryoEM.

### Negative stain EM

Purified Umb1 particles were diluted to 0.01 mg/mL and immediately subject to adsorption to glow-discharged carbon-coated copper grids for 60 seconds followed by 2% uranyl formate staining. Micrographs were recorded using Leginon^43^ on a 120 KV FEI Tecnai G2 Spirit with a Gatan Ultrascan 4000 4k x 4k CCD camera at 67,000 nominal magnification. The defocus ranged from -1.0 to -2.0 µm and the pixel size was 1.6 Å. The parameters of the contrast transfer function (CTF) were estimated using CTFFIND^44^. All particles were picked in a reference-free manner using DoG Picker^45^. The particle stack from the micrographs was pre-processed in Relion^46^. Particles were re-extracted with a binning factor of 4, resulting in a final box size of 80 pixels and a final pixel size of 6.4 Å. The reference-free 2D classification were performed using CryoSPARC^47^.

### CryoEM sample preparation, data collection and data processing

3 μL of 3 mg/mL purified Umb1 particle samples were loaded onto freshly glow discharged R 2/2 UltrAuFoil grids prior to plunge freezing using a Vitrobot Mark IV (ThermoFisher Scientific) with a blot force of 0 and 6 sec blot time at 100% humidity and 22°C. The data were acquired using an FEI Titan Krios transmission electron microscope operated at 300 kV and equipped with a Gatan K3 direct detector and Gatan Quantum GIF energy filter, operated in zero-loss mode with a slit width of 20 eV. Automated data collection was carried out using Leginon at a nominal magnification of 105,000× with a pixel size of 0.843 Å. 16,793 micrographs were collected with a defocus range comprised between -0.5 and -2.5 μm, respectively. The dose rate was adjusted to 15 counts/pixel/sec, and each movie was acquired in super-resolution mode fractionated in 75 frames of 40 ms. Movie frame alignment, estimation of the microscope contrast-transfer function parameters, particle picking, and extraction were carried out using Warp^48^. Two rounds of reference-free 2D classification were performed using CryoSPARC^47^ to select well-defined particle images. These selected particles were subjected to two rounds of 3D classification with 50 iterations each (angular sampling 7.5° for 25 iterations and 1.8° with local search for 25 iterations) using Relion^46^ with an initial model generated with ab-initio reconstruction in CryoSPARC. 3D refinements were carried out using non-uniform refinement along with per-particle defocus refinement in CryoSPARC. Selected particle images were subjected to the Bayesian polishing procedure^49^ implemented in Relion 3.1 before performing another round of non-uniform refinement in CryoSPARC followed by per-particle defocus refinement and again non-uniform refinement. Reported resolutions are based on the gold-standard Fourier shell correlation (FSC) of 0.143 criterion and Fourier shell correlation curves were corrected for the effects of soft masking by high-resolution noise substitution^50,51^.

### Umb1 particle model building and refinement

An initial structural model for the Umb1 particle was generated by combining the AlphaFold2 prediction of UmbC1 with the individual ALF-repeat:UmbB models and the UmbB:UmbA models. First, an UmbC1 model was docked into density and refined with cryoEM restraints^52^. This was then used as a reference model to align five individual ALF-repeat:UmbB models. Finally, the resulting UmbC1:5×UmbB1 model was used as a reference model to align the UmbB:UmbA models. It was not possible to determine the identities of individual UmbA subunits due to poor density and probable heterogeneity of the particle in the data. Therefore, two copies of UmbA1 and one each of UmbA4, UmbA5, and UmbA6 were included in the model, reflecting the relative abundance of these proteins in IP-MS analysis of proteins that interact with UmbC1. The 11-subunit model showed moderate agreement to density; the UmbB subunits matched reasonably well, but the orientations of the UmbB:UmbA interfaces and the UmbA domains were inconsistent. After refining to the density, a final model with good density agreement was produced. Using *density_tools* in Rosetta^52^, we calculated model/map FSC curves, which reveal a 0.5 crossing at 6.8 Å resolution.

### UmbC toxicity analysis in *S. aureus*

For analysis of the toxicity of UmbC toxin domains in a heterologous host, the xylose inducible plasmid pEPSA5 harboring the toxin of interest or empty vector were miniprepped from *S. aureus* and transformed in technical triplicate into competent RN4220 by electroporation, followed by one hour recovery in B2 medium at 37℃ 220 r.p.m. Transformations were plated on LBG supplemented with chloramphenicol and 0.2% w/v xylose to induce toxin expression. Transformant colonies were enumerated, and transformation efficiencies of empty plasmid and toxin-containing plasmid were computed and compared.

### Mutational profiling of *E. coli* expressing the toxin domain of UmbC1

Three *E. coli* strains – MG1655 Δ*ung* pPSV39-CV-*umbD1* pSCrhaB2-*umbC1*, MG1655 Δ*ung* pPSV39-CV-*umbD1* pSCrhaB2(no insert) and MG1655 Δ*ung* pPSV39-CV-*dddAI* and pSCrhaB2-*dddA* (32641830)– were grown in overnight cultures in LB supplemented with 15 μg/ml gentamycin, 50 μg/ml trimethoprim and 160 μM IPTG. The cultures were diluted 1:100 into fresh medium without IPTG, incubated until OD_600_ = 0.6, then supplemented with 0.2% rhamnose for toxin induction. Genomic DNA was isolated from the cultures after 60 min of induction, sequencing libraries were prepared as described^53^ and sequenced on an Illumina iSeq. SNV profiling was performed using described analysis methods^53,54^.

### Preparation of concentrated supernatant for use in screening for Umb targets

Spores of *S. coelicolor* wild-type and *Δumb* derivative strains were inoculated in R5 medium and grown for 36 hr. The cultures were then back diluted 1:200 in 50 mL R5 medium for a total combined culture volume of 150 mL and incubated 24-30 hr until reaching OD_600_ ∼4 Cells were then pelleted by centrifugation at 21,000 x g for 30 min. The resulting supernatant was filtered with a 0.45 μm PES membrane vacuum filter then concentrated using 100 kDa cutoff Amicon concentrators until reaching a final volume of 3 mL. The concentrated supernatant was run over an Econo-Pac 10DG desalting column (Bio-Rad), aliquoted, and stored at -80℃ until use.

### Isolation of bacteria from soil used in Umb toxicity screening

Soil isolate strains used in the broad Umb sensitivity screen were collected from sorghum plants grown at the University of California’s Agriculture and Natural Resources Kearney Agriculture Research and Extension Center in Parlier, CA, as described previously^55,56^. Root samples were obtained from mature sorghum plants that had been subjected to a prolonged pre-flowering drought. Immediately after extraction of plants from the soil, roots were removed and placed in 25% glycerol for 30 mins, then placed on dry ice until they were transferred to -80°C. To remove soil, roots were placed in a phosphate buffer and sonicated briefly. They were subsequently vortexed for 60 sec in 99% ethanol, 6 mins in 3% NaOCl, and 30 sec in 99% ethanol to sterilize the root surface. Roots were washed twice in sterilized dH_2_O, and 100 μL of rinse water was plated to check surface sterility. Roots were then cut into 1 cm pieces and placed into 2 mL tubes with 25% glycerol and incubated for 30 mins at room temperature before storing at -80°C. One 2 mL tube of roots (approximately 200 mg) was thawed and placed in a sterile ceramic mortar with 1 mL PBS buffer. Root tissue was ground gently, to release endophytic bacteria into the solution while minimizing lysis of bacterial cells. The solution was serially diluted, and 100 μL dilutions (10^-1^, 10^-2^, and 10^-3^) were plated onto various media types: ISP2, M9 minimal media, Skim Milk, Tap Water Yeast Extract, and Humic Acid. Plates were placed at 30°C and growth was monitored daily. When colonies were visible, they were picked and streaked onto a fresh plate of ISP2, followed by subsequent streaks if necessary to eliminate contamination, until only a single morphology was observed. The 16S ribosomal V3-V4 RNA sequences of the isolates were determined by Sanger sequencing.

### Screening diverse organisms for sensitivity to *S. coelicolor* Umb toxins

Strains used in this assay included both isolates obtained from culture collections, and a subset isolated in this study from the rhizosphere of field-grown sorghum plants (see above); all strains used in the assay and their growth conditions are listed in Table S6. Strains were grown as described at 30℃. Optical densities of initial cultures for all bacteria were measured and used to prepare 1 mL samples at an OD_600_ of 0.01 in the appropriate medium for each strain. 90 uL of each sample were transferred in duplicate to adjacent wells in a 96-well plate. To one of these wells, 10 uL of Umb supernatant from *S. coelicolor* was added. To the other well, 10 uL of *Δumb* supernatant from *S. coelicolor Δumb* was added. The plates were then incubated in a BioTek LogPhase 600 Microbiology Reader set to incubate the plates at 30℃ shaking at 800 r.p.m. taking OD_600_ measurements every 20 minutes for a total of 48 hr. Growth curves were monitored for the beginning of exponential phase. When an organism reached the beginning of its exponential growth phase, the corresponding duplicate cultures were removed from the incubator, combined with 100 uL BacTiter-Glo Reagent (Promega BacTiter-Glo™ Microbial Cell Viability Assay), and incubated at room temperature for 7 minutes. The luminescent signal was measured in a BioTek Cytation 1 imaging reader.

### Validation of initial hits from diverse organism Umb sensitivity assay

Potential target strains *S. griseus* NRRL B-2682, *S. ambofaciens* SAI 080, and *S. ambofaciens* SAI 195 along with negative control strain *S. mobaraensis* NRRL B-3729 were grown on SFM plates for three days. Colonies from these plates were excised and used to inoculate 30 mL TSBY and incubated 20 hr (*S. ambofaciens* and *S. griseus*) or 36 hr (*S. mobaraensis*) before being prepared for the Umb supernatant sensitivity assay as described above. Assay plates were initially incubated in the LogPhase for 7 hours. Samples were then collected, combined with BacTiter-Glo Reagent, and luminescence measured approximately hourly for a total of nine time points until the plates reached 20 hours total growth. At 16 hr, samples of each culture were serially diluted and plated on ISP2 agar to obtain an independent measure of growth yield.

### Streptomyces co-culture competition assays

For growth competition experiments between *Streptomyces* species, *S. coelicolor* spores were first inoculated into two 50 mL TSBY cultures and grown for ∼36 hr. Apramycin resistant *S. griseus* was similarly inoculated in TSBY and grown for 20 hr. When *S. coelicolor* cultures reached an OD_600_ = 3, 10 mL was aliquoted into four replicate baffled flasks. *S. griseus* cells washed twice with TSBY were then added to the culture flasks at OD_600_ = 0.03, establishing an initial *S. coelicolor: S. griseus* ratio of 100:1. Cultures were serially diluted and plated on selective (for *S. griseus*) and nonselective media (total population) for c.f.u. quantification at an initial time point and after incubation at 28℃ for 12 hours.

### Microscopy

Imaging was performed on a Nikon Eclipse Ti-E wide-field microscope equipped with a sCMOS camera (Hamamatsu). A 60X 1.4 NA oil-immersion PH3 objective was used for imaging. The microscope was controlled by NIS-Elements v3.30.02. The microscope chamber was heated to 28°C, and *S. griseus* spores were loaded into all four chambers of a bacterial microfluidic plate (B04 from EMD Millipore). Using a CellASIC ONIX (Model EV262) microfluidic perfusion system, a pressure of 2 PSI was applied to two columns over two roughly 6 hr intervals. One chamber was treated with media and Umb supernatant for interval one (0-350 min) followed by media alone for interval two (350-660 min). A second chamber was treated with media and *Δumb* supernatant followed by media alone. A third chamber was treated with media alone followed by media and Umb supernatant. Finally, a fourth chamber was treated with PBS followed by media alone.

Z stacks were acquired at each of 3 positions in each imaging chamber every 10 min. Z stacks were merged using gaussian focus stacking followed by automatic frame alignment in FIJI^57^. Cells that were imaged without occlusion or growth outside the field of view for the duration of 11 hr were manually selected and exported in Napari (doi:10.5281/zenodo.3555620) using the napari-crop and napari-nd-cropper plugins. Cells were automatically segmented frame-by-frame using Omnipose (bact_phase_omni model)^58^. Spurious labels arising from plate defects, debris, or pillars were removed manually in Napari following automatic edge-based filtering in Python. Finally, cells were tracked (and any over-segmentation resolved) by manually recoloring Z stack labels in Napari using the fill tool in 3D mode. All processed spacetime labels were then loaded into Python for extracting area over time per cell.

### Bioinformatics analysis

To comprehensively retrieve UmbC protein homologs, the PSI-BLAST program^59^ was employed for iterative searches against the NCBI non-redundant (nr) protein database until convergence, with a cut-off e-value of 0.005. The five upstream and five downstream gene neighbors of UmbC were extracted from the NCBI GenBank files for use in the gene neighborhood analysis^60^. All protein neighbors were clustered based on their sequence similarities using the BLASTCLUST program, a BLAST score-based single-linkage clustering method (https://ftp.ncbi.nih.gov/blast/documents/blastclust.html). Protein clusters were then annotated based on their domain architectures using the HMMSCAN program^61^, searching against the Pfam database^62^ and our in house custom HMM profile database. Signal peptide and transmembrane region prediction was determined using the Phobius program^63^. For systematic identification and classification of C-terminal toxin domains in UmbC proteins and the immunity families represented by UmbD proteins, we utilized the CLANS program (https://doi.org/10.1093/bioinformatics/bth444). This program employed a network analysis to organize sequences through the application of the Fruchterman and Reingold force-directed layout algorithm (https://doi.org/10.1002/spe.4380211102) based on their sequence similarities derived from all-against-all BLASTP comparisons. A representative sequence of the novel domain family served as a seed in PSIBLAST searches to retrieve homologs. Following removal of highly similar sequences by BLASTCLUST, multiple sequence alignments (MSA) were built using KALIGN^64^, MUSCLE^65^ or PROMALS3D^66^. To identify the conserved residues for each domain family, a custom Perl script was used to calculate the conservation pattern of the MSA based on different categories of amino acid physio-chemical properties developed by Taylor^67^. Structural models for representative sequences of each domain family were predicted using AlphaFold2^17^ and models with the highest predicted Local Distance Difference Test (LDDT) scores were selected. Determination of domain boundaries for each family was guided by both the structure models and the PAE matrix provided by AlphaFold2. Functional predictions for toxin domains belonging to uncharacterized families were generated using DALI^68^and Foldseek^69^ searches with representative structural models from each family to identify structurally-related proteins with characterized functions. Function predictions were assigned when structurally similar proteins or protein domains (DALI z score >3, or Foldseek E-value <0.01) with known toxin activities were identified.

## Acknowledgements

We thank Simon Dove, Joshua Woodward, E. Peter Greenberg and Carrie Harwood for helpful discussions, Ricard Rodriguez for help with running and troubleshooting mass spectrometry samples, Linquan Bai and Xinran Wang for providing the plasmids pSET152 and pKGLP2, respectively, Eoin Brodie for providing wheat rhizosphere isolates, and the USDA-ARS Culture Collection (NRRL) for providing strains. This study was supported by Defense Advanced Research Projects Agency Biological Technologies Office Program: Harnessing Enzymatic Activity for Lifesaving Remedies (HEALR) under cooperative agreement no. HR0011-21-2-0012 (to J.D.M.), the National Institute of Allergy and Infectious Diseases (75N93022C00036 to D.V.), a Pew Biomedical Scholars Award (D.V.), an Investigators in the Pathogenesis of Infectious Disease Awards from the Burroughs Wellcome Fund (D.V), the University of Washington Arnold and Mabel Beckman cryoEM center, National Institute of Health grant S10OD032290 (to D.V.), a Saint Louis University Startup Fund (to D.Z.), the US Department of Agriculture (CRIS 2030-21430-008-00D to D.C.D.), and USDA-NIFA (2019-67019-29306 to D.C.D), and is a contribution of the Pacific Northwest National Laboratory (PNNL) Secure Biosystems Design Science Focus Area “Persistence Control of Engineered Functions in Complex Soil Microbiomes” (operated by the U.S. DOE under contract DE-AC05-76RL01830 to D.C.D.). J.D.M. and D.V. are HHMI Investigators, D.V. and J.D.M hold the Hans Neurath Endowed Chair in Biochemistry and the Lynn M. and Michael D. Garvey Endowed Chair in Gastroenterology, respectively, at the University of Washington.

## Supplemental Figure Legends

**Video S1.** Time lapse microscopy analysis of *S. griseus* cells undergoing Umb-mediated intoxication. Cells were exposed to the indicated treatments during growth in a flow cell; rectangular objects are flow cell structures.

---

**Figure S1:**
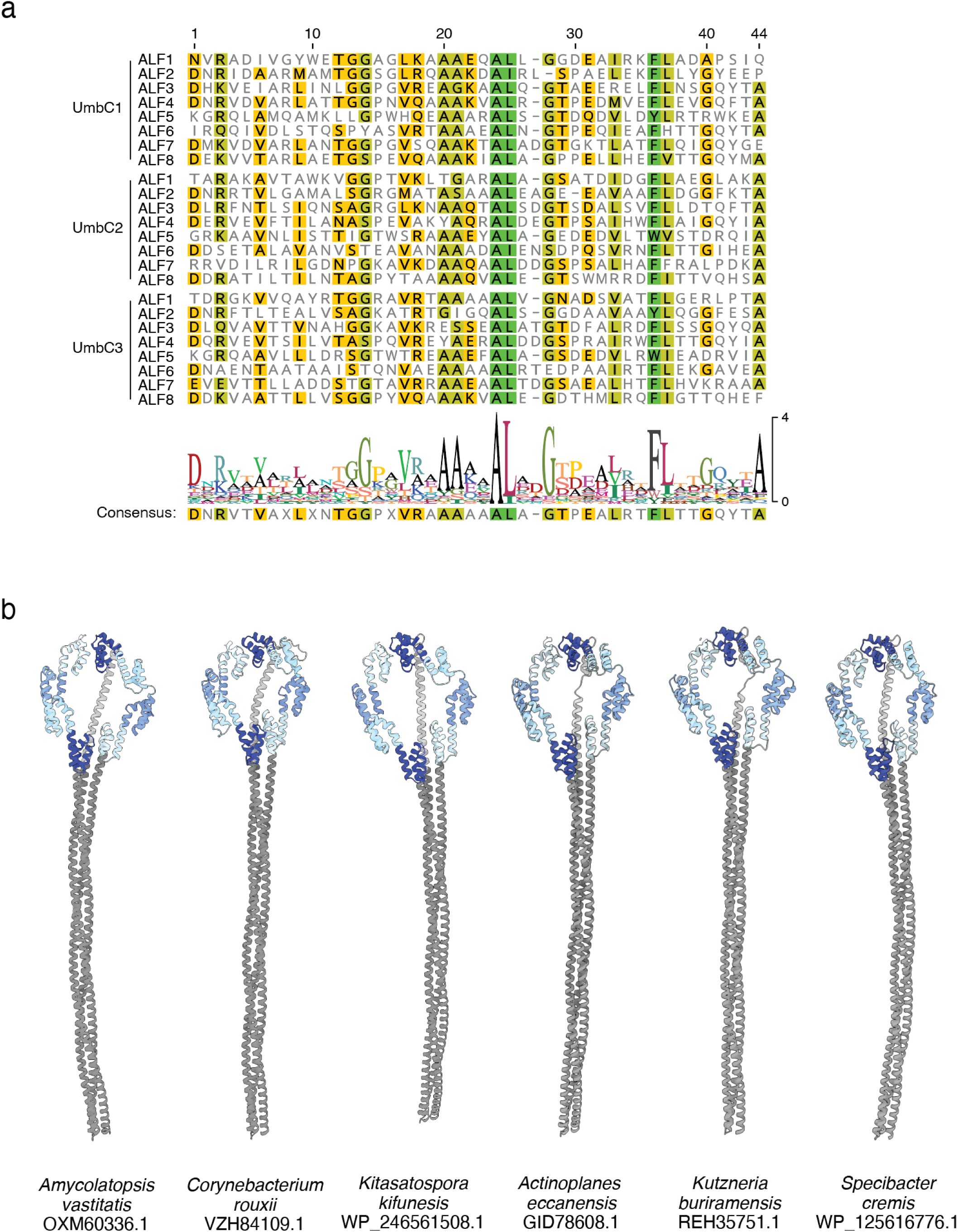
Degenerate nature of ALF repeat sequences and example UmbC structural models with straight coiled-coil domains. **a,** Alignment of ALF repeats 1-8 from each UmbC protein of *S. coelicolor.* The minimum ALF repeat unit was selected based on the structural model. **b,** Predicted structural models of assorted UmbC proteins, obtained using default AlphaFold parameters and without templating.

**Figure S2:**
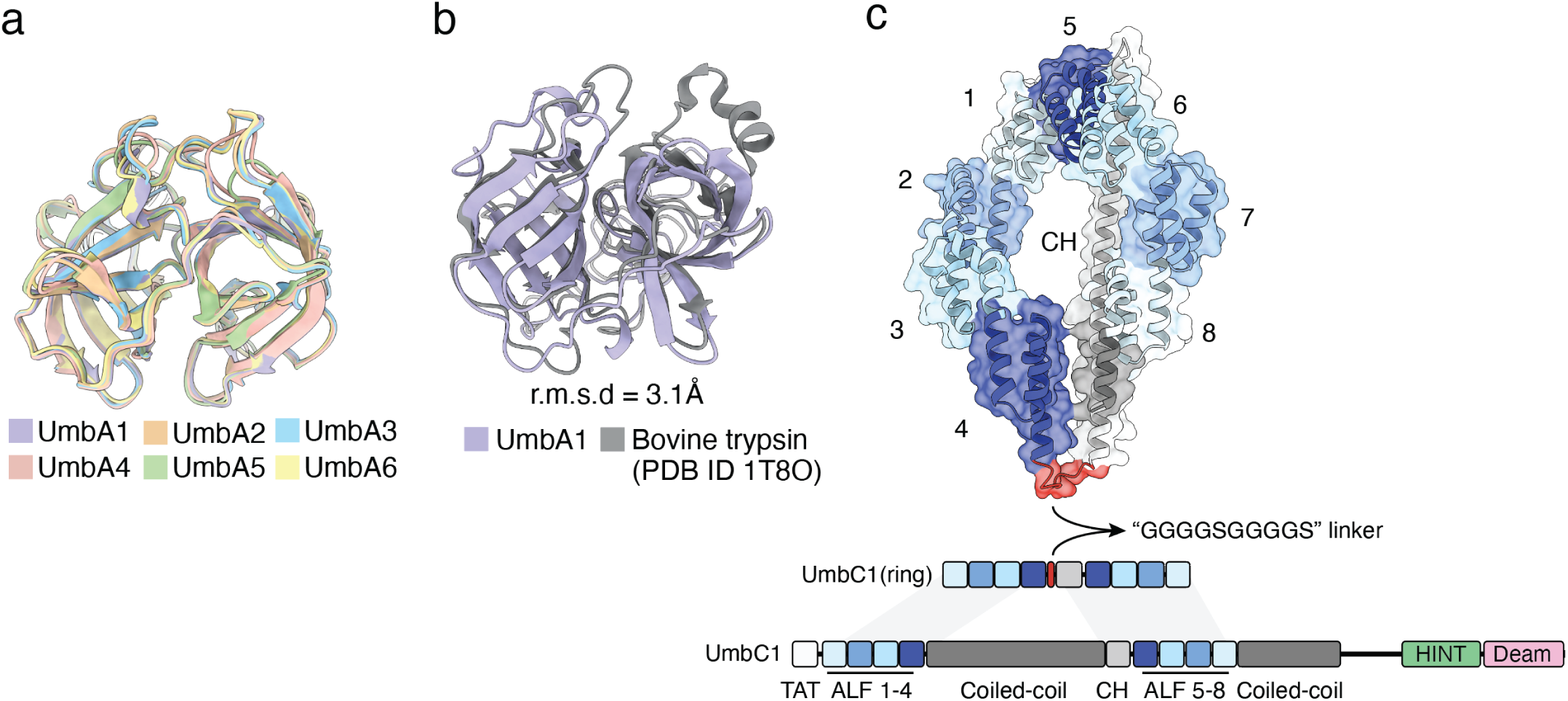
UmbA proteins contain a conserved trypsin-like domain, and design of a construct for the expression of UmbC1(ring). **a,** Alignment of the trypsin-like domain of the UmbA proteins of *S. coelicolor.* Numbers indicate amino acid positions included; signal sequences were removed for clarity. **b,** Alignment of UmbA1(T) and bovine trypsin. **c**, Predicted structure and genetic architecture of our construct for the expression of UmbC1(ring).

**Figure S3:**
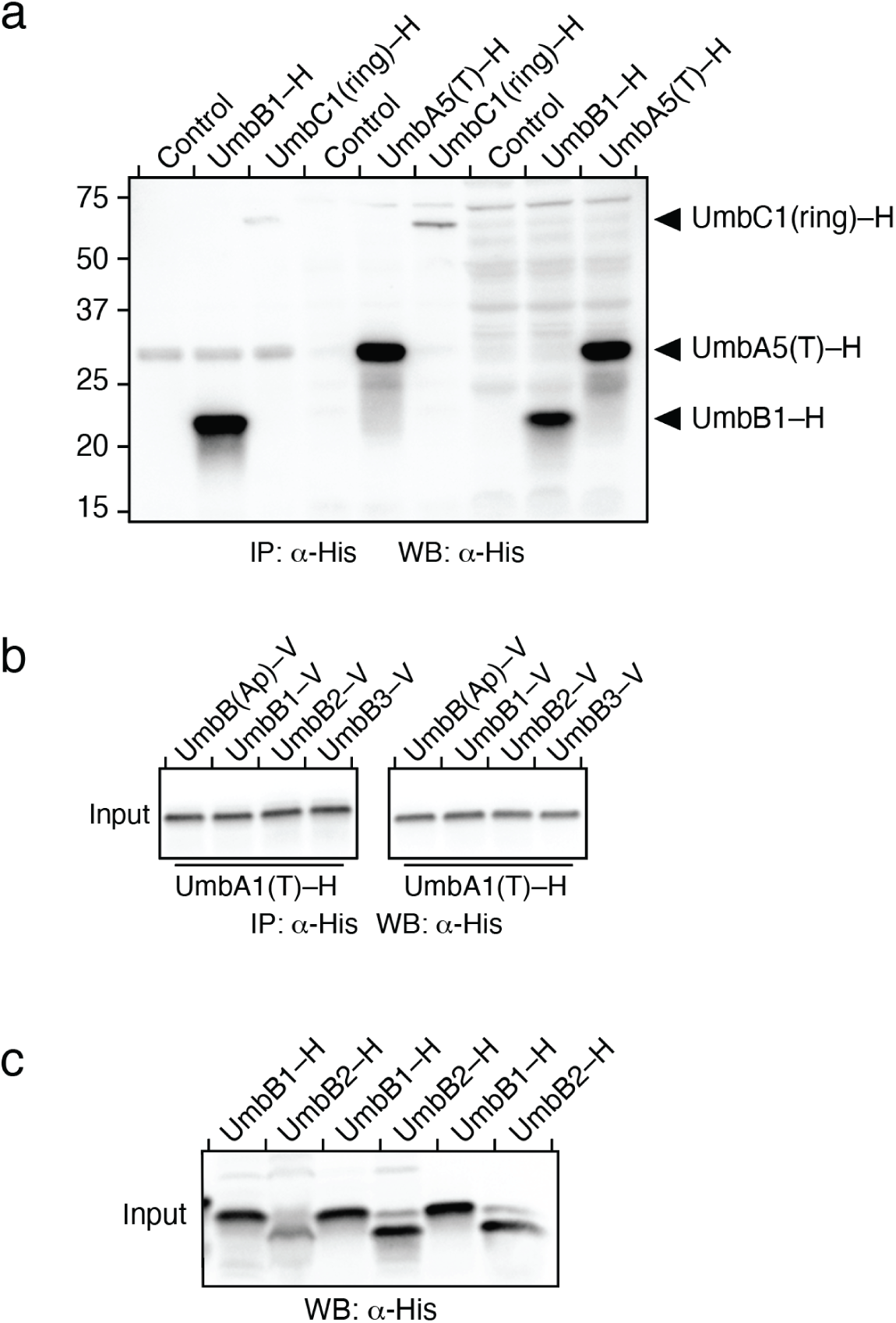
Input protein levels from studies of the interactions between proteins in the Umb complex. **a-c**, WB analyses of input samples from IP experiments between the indicated heterologously expressed, tagged (–H, hexahistidine; –V, VSV-G epitope) Umb proteins. Controls lanes correspond to beads in the absence of a bait protein. UmbB(Ap) is a UmbB protein from the distantly related species *Actinoplanes philippinensis*.

**Figure S4:**
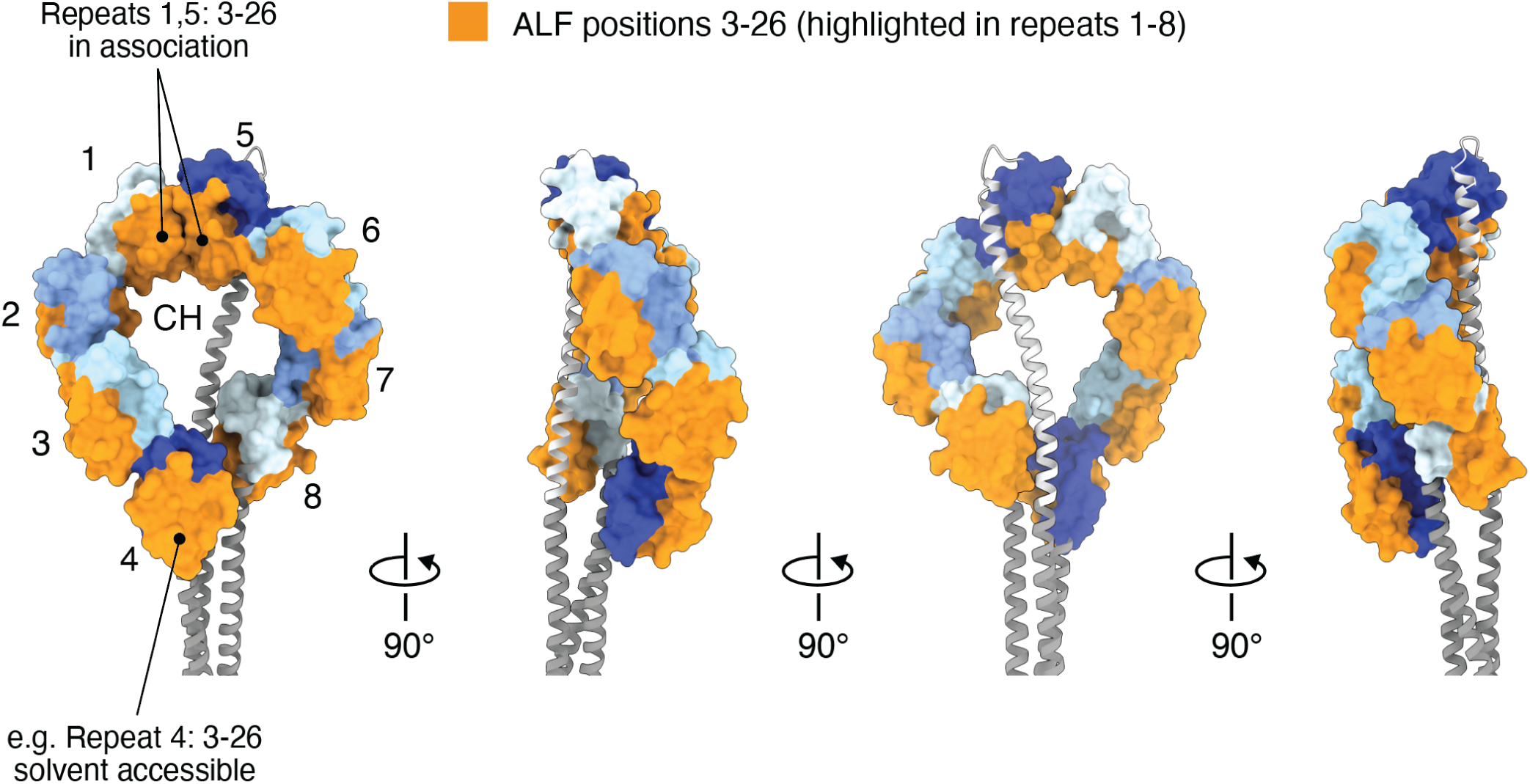
ALF repeats 1 and 5 exhibit a distinct orientation. Orange coloring indicates the residues of the ALF repeats of UmbC1 that are exposed to the surface in repeats predicted to interact with AtrB in structural models (ALF 2,3, 4-8). In repeats 1 and 5, many of these residues are buried in the interface between the ALF repeats.

**Figure S5:**
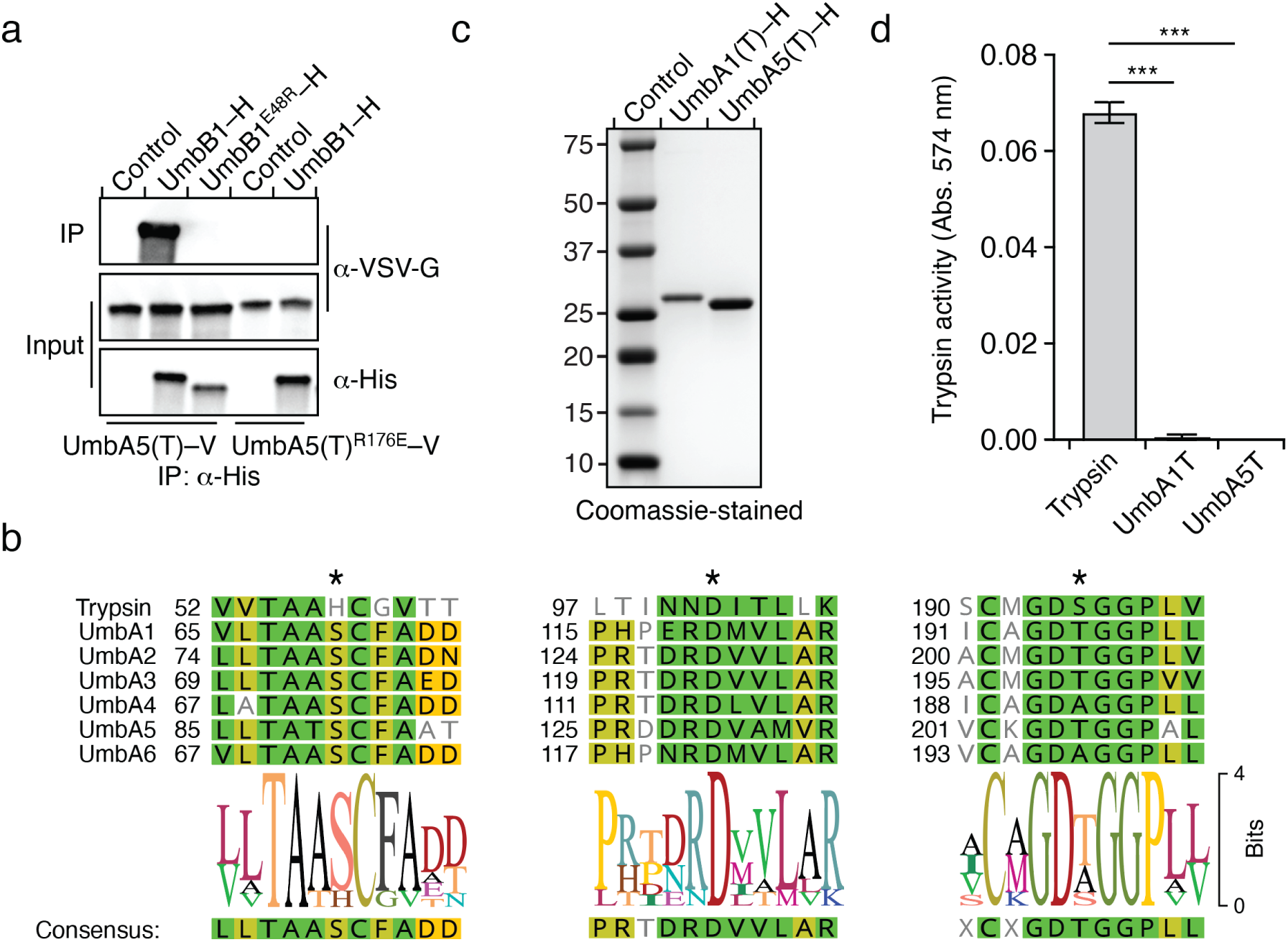
The trypsin-like domain of UmbA proteins mediates binding with UmbB and lacks catalytic activity. **a,** WB analysis from IP experiments of the indicated heterologously expressed, tagged Umb proteins. UmB1^E48R^–H and UmbA5(T)^R176E^–V contain substitutions of residues predicted to be critical for interaction between the two proteins. **b,** Structure-guided alignments of the UmbA(T) regions normally encompassing the catalytic histidine, aspartate, serine triad typical of trypsin proteins, indicating the conserved substitutions found across the UmbA proteins of *S. coelicolor.* **c, d,** Coomassie-stained SDS-PAGE analysis (c) and proteolytic activity (d) of purified, heterologously expressed UmbA1(T) and UmbA5(T).

**Figure S6:**
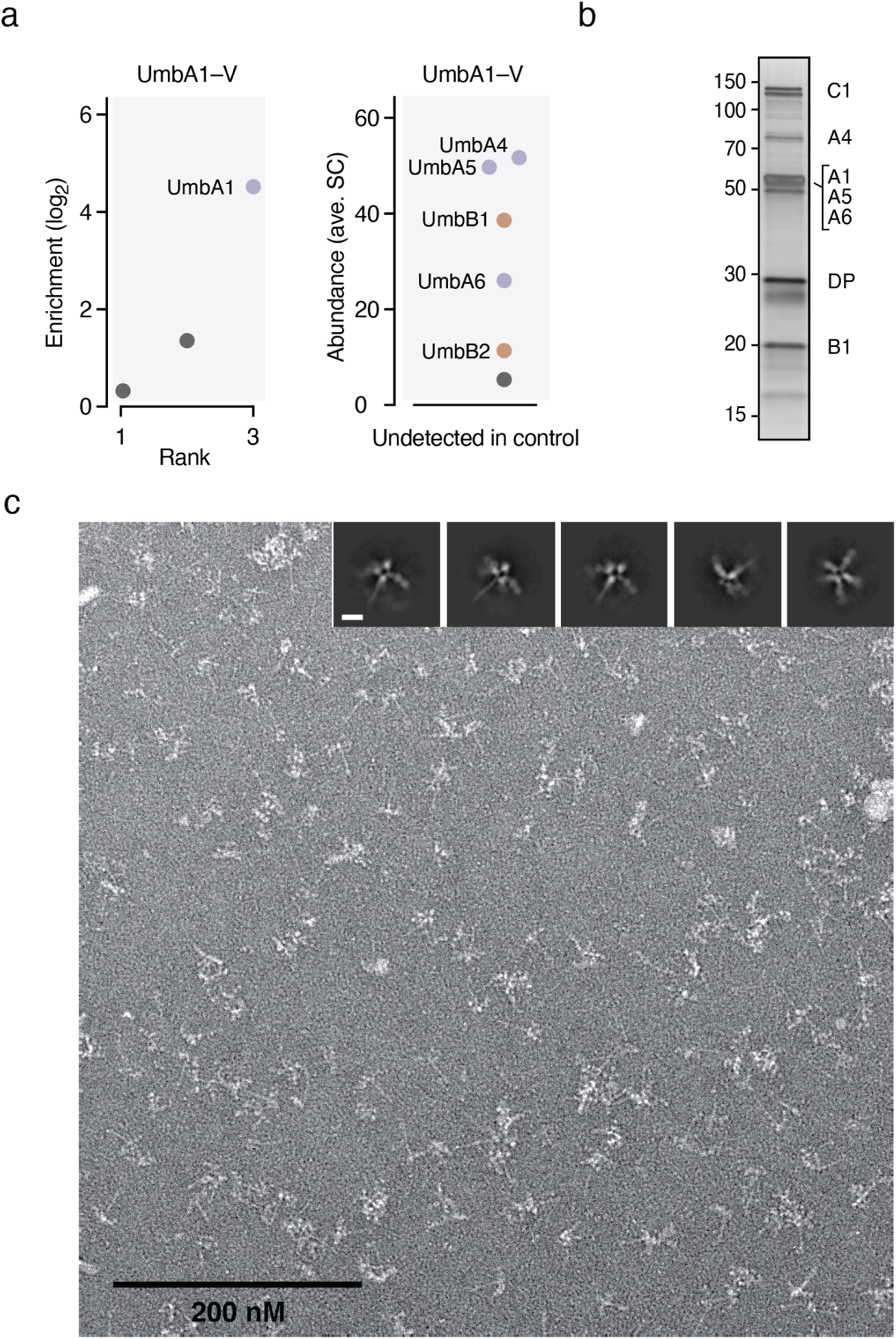
Purification of the Umb1 complex using epitope-tagged UmbA1 yields a protein complex with an umbrella-like morphology. **a,** IP-MS identification of proteins that interact with UmbA1-VSV-G. Left panel indicates the average fold enrichment of proteins detected in both IP and control samples; right panel presents abundance (average spectral counts, SC) for proteins detected only in IP samples. Colors indicate paralogous proteins and correspond to Figure 2; non-Umb proteins shown in grey. n = 2 biological replicates. **b,** Silver-stained SDS-PAGE analysis of the Umb1 particle, purified using UmbA1–8xHis, with bands corresponding to individual Umb1 proteins identified. DP, degradation product. **c**, Full field view of TEM analysis of negative stained, purified Umb1 particles. Insets show a selection of class averages with particles adopting different orientations. Inset scale bar, 100 Å.

**Figure S7:**
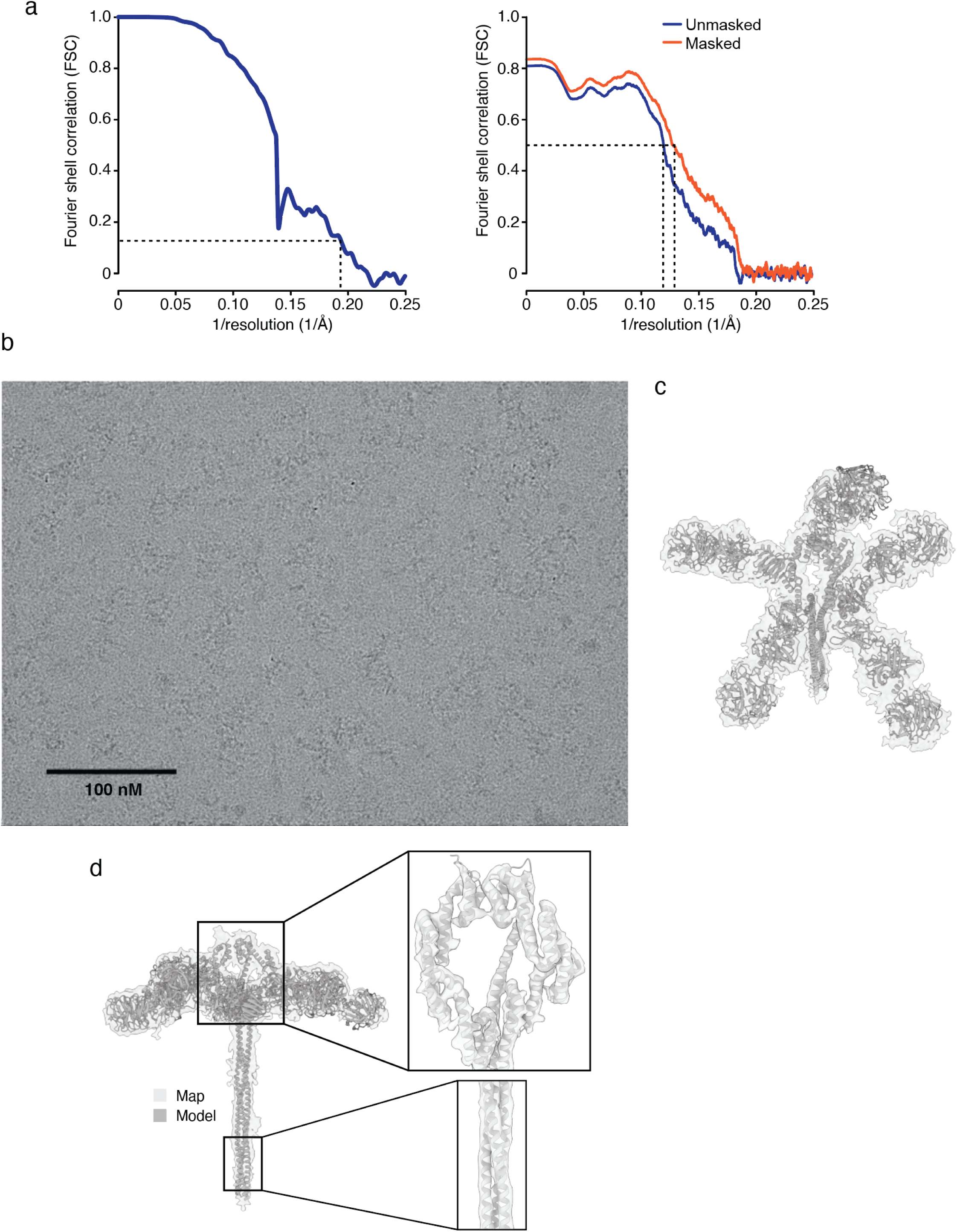
Cryo-EM based structural characterization of the Umb1 particle. **a,** Masked and unmasked map vs. model FSC plot for the Umb1 particle. Map resolution is 5.1 Å using the gold-standard FSC cut-off of 0.143 (left) and is 6.8 Å using map versus model at an FSC cut-off of 0.5 (right). **b,** Representative micrograph from the UmbC1 cryo-EM dataset. **c,d,** Cryo-EM density corresponding to the full UmbC1 model. Insets in (d) show portions of the ring and stalk, highlighting the clarity of secondary structure in regions of our maps.

**Figure S8:**
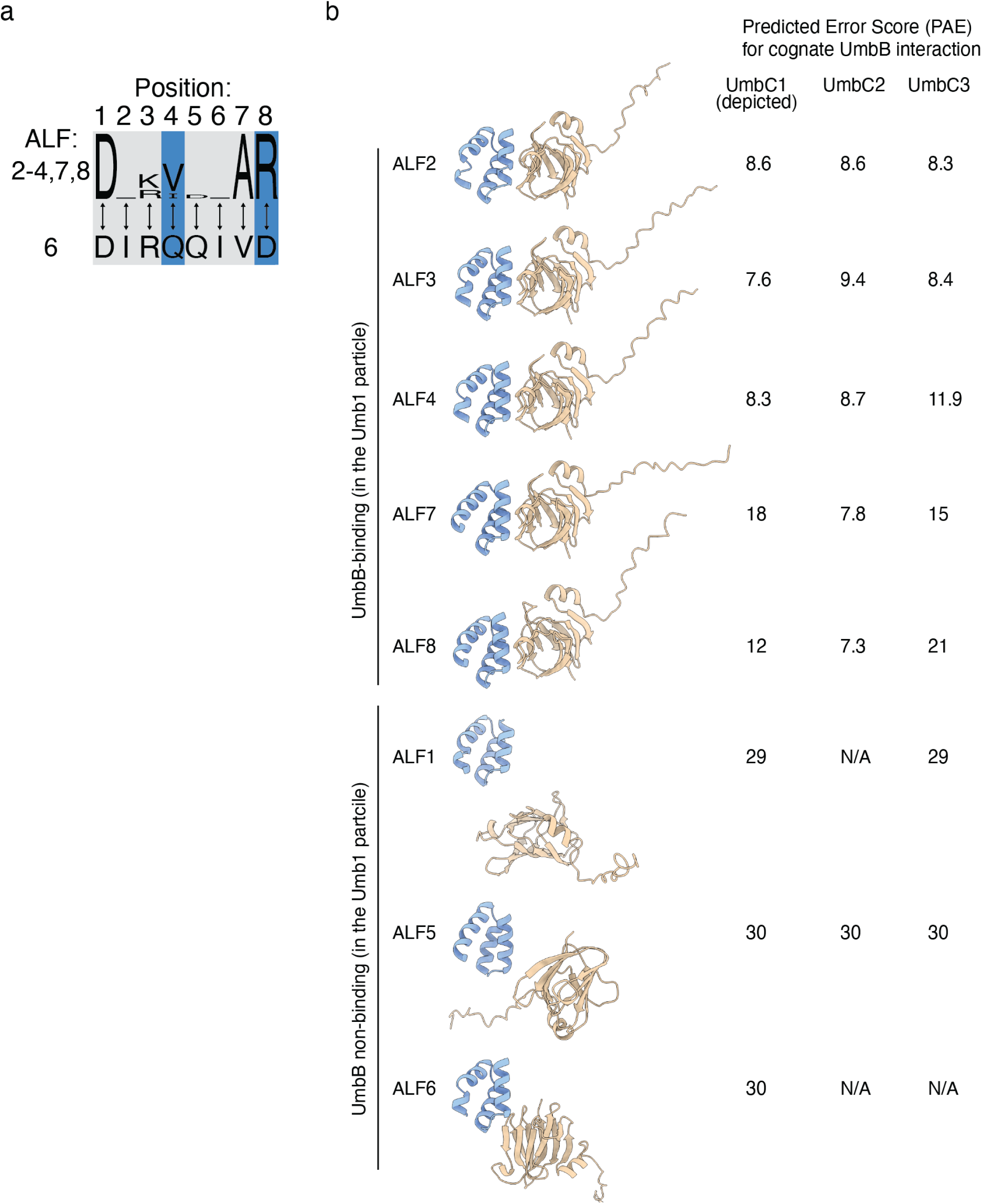
Structure and sequence-based differentiation of UmbB-interacting and non-interaction ALF repeats. **a,** Probability sequence logo generated from an alignment of positions 1-8 of the UmbB-interacting ALF repeats of *S. coelicolor,* compared to the analogous positions in AFL6. Positions located at the interaction interface and which have non-conservative substitutions in ALF6 are highlighted in blue. **b,** Predicted structural models for the interaction between each ALF repeat of UmbC1 with UmbB1, and RoseTTAFold2 predicted error scores (PAE) calculated for models of the ALF repeats of each *S. coelicolor* UmbC protein interacting with its cognate UmbB. PAE values: <10, high confidence; <20, moderate confidence; >20, low confidence^38^. N/A, no interaction predicted.

**Figure S9:**
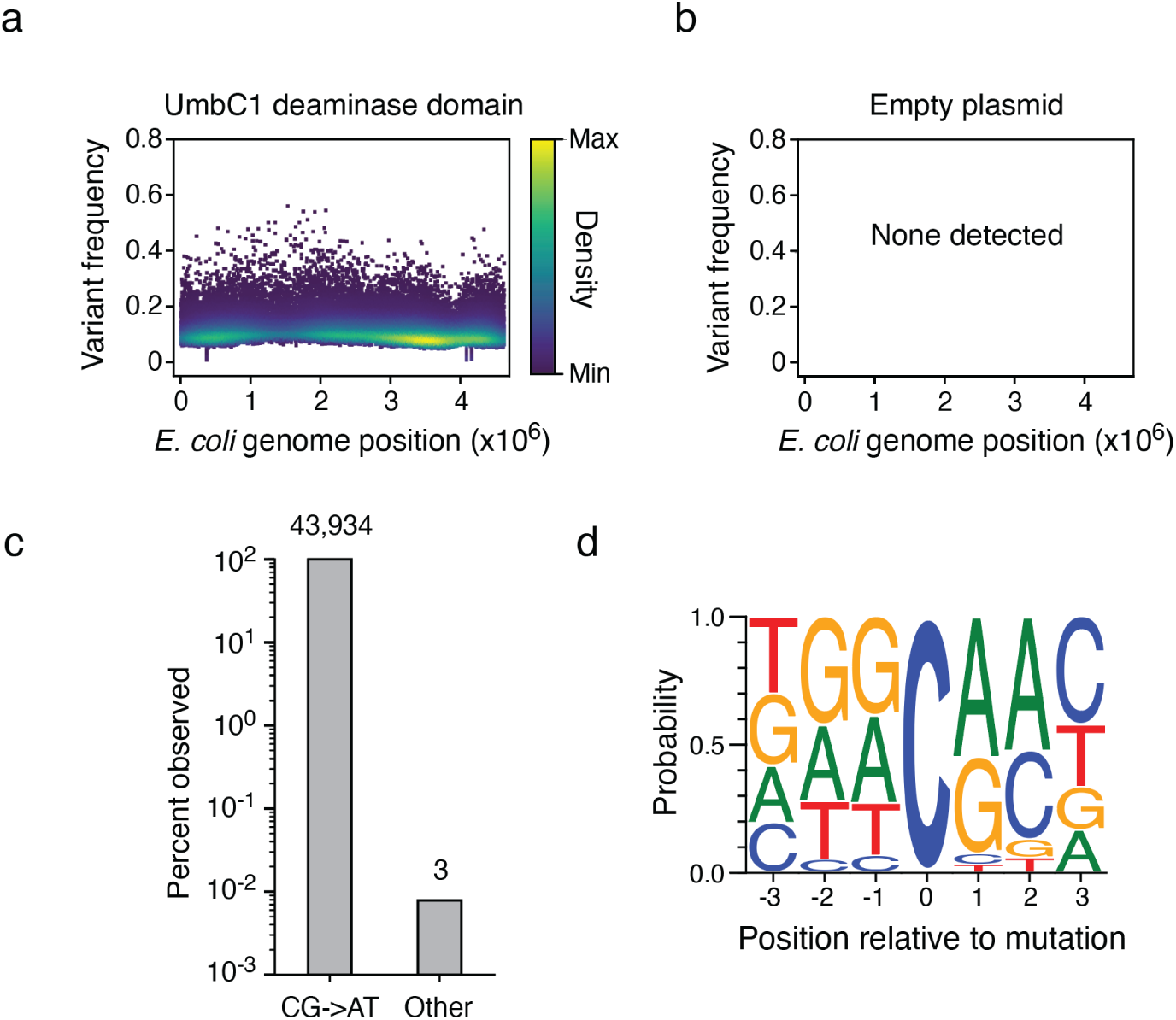
The toxin domain of UmbC1 exhibits mutagenic cytosine deaminase activity. **a,b,** Representation of single-nucleotide variants (SNVs) by chromosomal position, frequency, and density in *E. coli* Δ*ung* following 60 min induction of expression of the deaminase toxin domain from UmbC1 (a), or the equivalently-treated vector control strain (b). **c,** Frequency of the indicated substitutions among the SNVs shown in (a). **d,** Probability sequence logo of the region flanking mutated cytosines among the SNVs shown in (a).

**Figure S10:**
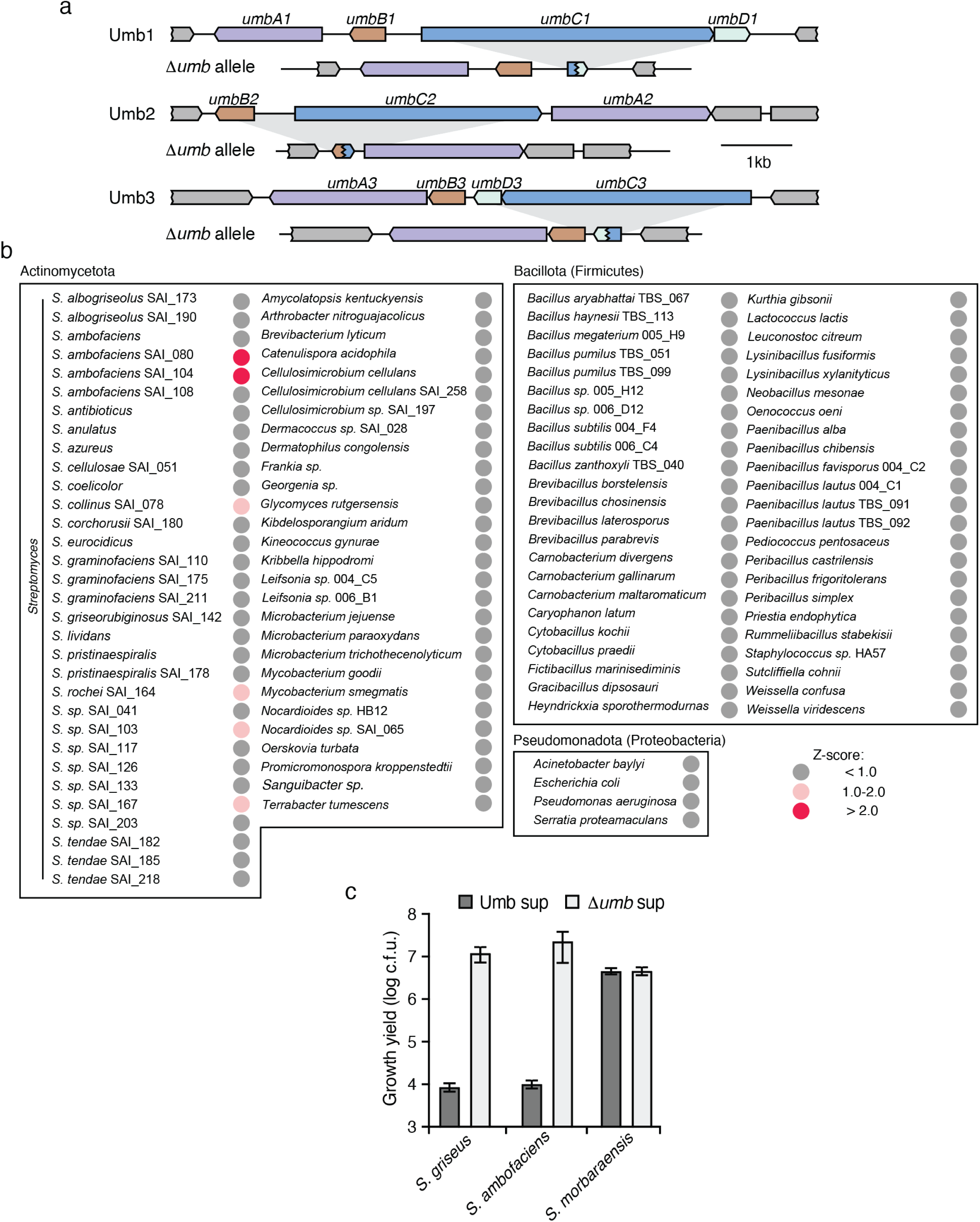
Screen of diverse soil bacteria to identify targets of the Umb toxins of *S. coelicolor.* **a,** Genetic loci schematic indicating deletions present in *S. coelicolor Δumb.* **b,** Umb toxin target screening results for strains not depicted in Figure 4b, grouped by target strain phylum. Z-scores were calculated as in Fig 4b; scores >2 indicate significant Umb-dependent inhibition. **c**, Growth yields (c.f.u, colony forming units) determined of the indicated strains grown in Umb or Δ*umb* supernatant for 16 hr.

**Figure S11:**
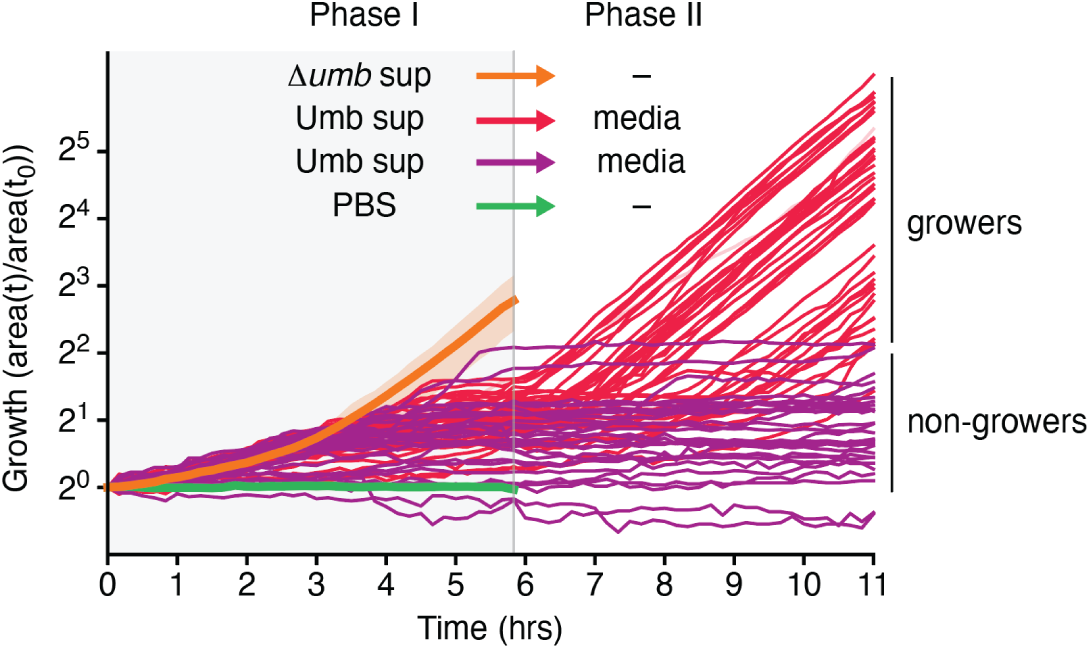
Growth trajectories of individual *S. coelicolor* cells treated with Umb supernatant. After exchange of Umb supernatant with fresh medium, a portion of the population resumes growth (growers) while other treated cells remain arrested (non-growers). Average growth of other treatment groups show only in Phase I for clarity.

**Figure S12:**
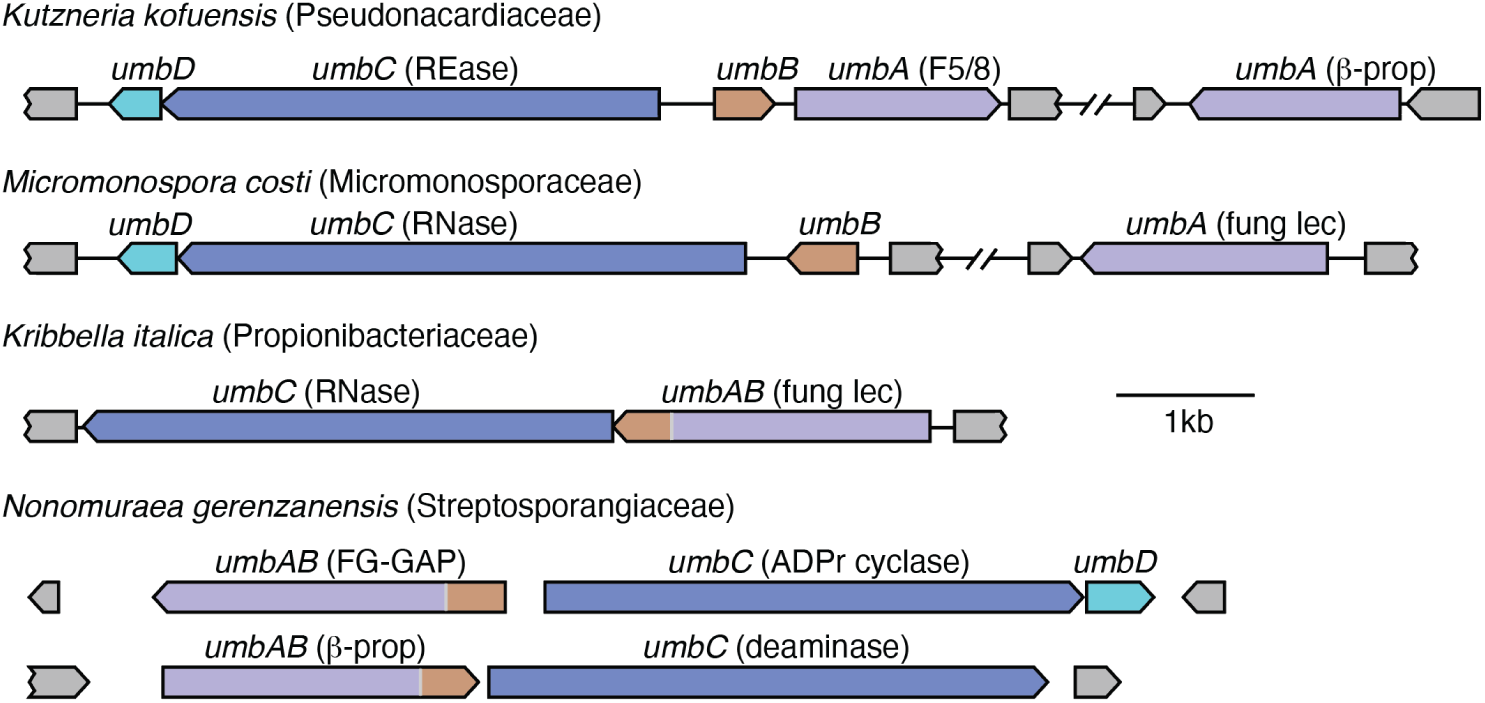
Representative *umb* loci from phylogenetically diverse Actinobacteria. Toxin domains encoded by the *umbC* genes and lectin domains encoded by the *umbA* genes are defined in Tables S2 and S4, respectively. β-prop, UAL-Bprop-1 family identified in this study.

## Bibliography

1. Anthony, M.A., Bender, S.F., and van der Heijden, M.G.A. (2023). Enumerating soil biodiversity. Proc Natl Acad Sci U S A 120, e2304663120. 10.1073/pnas.2304663120.

2. Granato, E.T., Meiller-Legrand, T.A., and Foster, K.R. (2019). The Evolution and Ecology of Bacterial Warfare. Curr Biol 29, R521–R537. 10.1016/j.cub.2019.04.024.

3. Peterson, S.B., Bertolli, S.K., and Mougous, J.D. (2020). The Central Role of Interbacterial Antagonism in Bacterial Life. Curr Biol 30, R1203–R1214. 10.1016/j.cub.2020.06.103.

4. Alam, K., Mazumder, A., Sikdar, S., Zhao, Y.M., Hao, J., Song, C., Wang, Y., Sarkar, R., Islam, S., Zhang, Y., and Li, A. (2022). Streptomyces: The biofactory of secondary metabolites. Front Microbiol 13, 968053. 10.3389/fmicb.2022.968053.

5. Barka, E.A., Vatsa, P., Sanchez, L., Gaveau-Vaillant, N., Jacquard, C., Meier-Kolthoff, J.P., Klenk, H.P., Clement, C., Ouhdouch, Y., and van Wezel, G.P. (2016). Taxonomy, Physiology, and Natural Products of Actinobacteria. Microbiol Mol Biol Rev 80, 1–43. 10.1128/MMBR.00019-15.

6. Hopwood, D.A. (2007). Streptomyces in Nature and Medicine (Oxford University Press).

7. Kinkel, L.L., Schlatter, D.C., Xiao, K., and Baines, A.D. (2014). Sympatric inhibition and niche differentiation suggest alternative coevolutionary trajectories among Streptomycetes. ISME J 8, 249–256. 10.1038/ismej.2013.175.

8. Aoki, S.K., Diner, E.J., de Roodenbeke, C.T., Burgess, B.R., Poole, S.J., Braaten, B.A., Jones, A.M., Webb, J.S., Hayes, C.S., Cotter, P.A., and Low, D.A. (2010). A widespread family of polymorphic contact-dependent toxin delivery systems in bacteria. Nature 468, 439–442. nature09490 [pii] 10.1038/nature09490.

9. Hood, R.D., Singh, P., Hsu, F., Guvener, T., Carl, M.A., Trinidad, R.R., Silverman, J.M., Ohlson, B.B., Hicks, K.G., Plemel, R.L., et al. (2010). A type VI secretion system of Pseudomonas aeruginosa targets a toxin to bacteria. Cell Host Microbe 7, 25–37.

10. Jamet, A., and Nassif, X. (2015). New players in the toxin field: polymorphic toxin systems in bacteria. MBio 6, e00285–00215. 10.1128/mBio.00285-15.

11. Klein, T.A., Ahmad, S., and Whitney, J.C. (2020). Contact-Dependent Interbacterial Antagonism Mediated by Protein Secretion Machines. Trends Microbiol 28, 387–400. 10.1016/j.tim.2020.01.003.

12. Souza, D.P., Oka, G.U., Alvarez-Martinez, C.E., Bisson-Filho, A.W., Dunger, G., Hobeika, L., Cavalcante, N.S., Alegria, M.C., Barbosa, L.R., Salinas, R.K., et al. (2015). Bacterial killing via a type IV secretion system. Nat Commun 6, 6453. 10.1038/ncomms7453.

13. Whitney, J.C., Peterson, S.B., Kim, J., Pazos, M., Verster, A.J., Radey, M.C., Kulasekara, H.D., Ching, M.Q., Bullen, N.P., Bryant, D., et al. (2017). A broadly distributed toxin family mediates contact-dependent antagonism between gram-positive bacteria. Elife 6, e26938. 10.7554/eLife.26938.

14. Zhang, D., de Souza, R.F., Anantharaman, V., Iyer, L.M., and Aravind, L. (2012). Polymorphic toxin systems: Comprehensive characterization of trafficking modes, processing, mechanisms of action, immunity and ecology using comparative genomics. Biol Direct 7, 18. 10.1186/1745-6150-7-18.

15. Ruhe, Z.C., Low, D.A., and Hayes, C.S. (2020). Polymorphic Toxins and Their Immunity Proteins: Diversity, Evolution, and Mechanisms of Delivery. Annu Rev Microbiol 74, 497–520. 10.1146/annurev-micro-020518-115638.

16. Yeats, C., Bentley, S., and Bateman, A. (2003). New knowledge from old: in silico discovery of novel protein domains in Streptomyces coelicolor. BMC microbiology 3, 3. 10.1186/1471-2180-3-3.

17. Jumper, J., Evans, R., Pritzel, A., Green, T., Figurnov, M., Ronneberger, O., Tunyasuvunakool, K., Bates, R., Zidek, A., Potapenko, A., et al. (2021). Highly accurate protein structure prediction with AlphaFold. Nature 596, 583–589. 10.1038/s41586-021-03819-2.

18. Rini, J.M. (1995). Lectin structure. Annu Rev Biophys Biomol Struct 24, 551–577. 10.1146/annurev.bb.24.060195.003003.

19. Hedstrom, L. (2002). Serine protease mechanism and specificity. Chem Rev 102, 4501–4524. 10.1021/cr000033x.

20. Dassa, B., Haviv, H., Amitai, G., and Pietrokovski, S. (2004). Protein splicing and auto-cleavage of bacterial intein-like domains lacking a C’-flanking nucleophilic residue. J Biol Chem 279, 32001–32007. 10.1074/jbc.M404562200.

21. Jeong, Y., Kim, J.N., Kim, M.W., Bucca, G., Cho, S., Yoon, Y.J., Kim, B.G., Roe, J.H., Kim, S.C., Smith, C.P., and Cho, B.K. (2016). The dynamic transcriptional and translational landscape of the model antibiotic producer Streptomyces coelicolor A3(2). Nat Commun 7, 11605. 10.1038/ncomms11605.

22. Kim, W., Hwang, S., Lee, N., Lee, Y., Cho, S., Palsson, B., and Cho, B.K. (2020). Transcriptome and translatome profiles of Streptomyces species in different growth phases. Sci Data 7, 138. 10.1038/s41597-020-0476-9.

23. Kwak, J., Jiang, H., and Kendrick, K.E. (2002). Transformation using in vivo and in vitro methylation in Streptomyces griseus. FEMS Microbiol Lett 209, 243–248. 10.1111/j.1574-6968.2002.tb11138.x.

24. Russel, J., Roder, H.L., Madsen, J.S., Burmolle, M., and Sorensen, S.J. (2017). Antagonism correlates with metabolic similarity in diverse bacteria. Proc Natl Acad Sci U S A 114, 10684–10688. 10.1073/pnas.1706016114.

25. Waksman, S.A. (1941). Antagonistic Relations of Microorganisms. Bacteriol Rev 5, 231–291.

26. Cascales, E., Buchanan, S.K., Duche, D., Kleanthous, C., Lloubes, R., Postle, K., Riley, M., Slatin, S., and Cavard, D. (2007). Colicin biology. Microbiol Mol Biol Rev 71, 158–229. 71/1/158 [pii] 10.1128/MMBR.00036-06.

27. Jakes, K.S., and Cramer, W.A. (2012). Border crossings: colicins and transporters. Annu Rev Genet 46, 209–231. 10.1146/annurev-genet-110711-155427.

28. Calcuttawala, F., Pal, A., Nath, P., Kar, R., Hazra, D., and Pal, R. (2021). Structural and functional insights into colicin: a new paradigm in drug discovery. Archives of microbiology 204, 37. 10.1007/s00203-021-02689-6.

29. Mavridou, D.A.I., Gonzalez, D., Kim, W., West, S.A., and Foster, K.R. (2018). Bacteria Use Collective Behavior to Generate Diverse Combat Strategies. Curr Biol 28, 345–355 e344. 10.1016/j.cub.2017.12.030.

30. Gygli, S.M., Borrell, S., Trauner, A., and Gagneux, S. (2017). Antimicrobial resistance in Mycobacterium tuberculosis: mechanistic and evolutionary perspectives. FEMS Microbiol Rev 41, 354–373. 10.1093/femsre/fux011.

31. Hennart, M., Panunzi, L.G., Rodrigues, C., Gaday, Q., Baines, S.L., Barros-Pinkelnig, M., Carmi-Leroy, A., Dazas, M., Wehenkel, A.M., Didelot, X., et al. (2020). Population genomics and antimicrobial resistance in Corynebacterium diphtheriae. Genome Med 12, 107. 10.1186/s13073-020-00805-7.

32. Myronovskyi, M., Welle, E., Fedorenko, V., and Luzhetskyy, A. (2011). Beta-glucuronidase as a sensitive and versatile reporter in actinomycetes. Applied and environmental microbiology 77, 5370–5383. 10.1128/AEM.00434-11.

33. Bierman, M., Logan, R., O’Brien, K., Seno, E.T., Rao, R.N., and Schoner, B.E. (1992). Plasmid cloning vectors for the conjugal transfer of DNA from Escherichia coli to Streptomyces spp. Gene 116, 43–49. 10.1016/0378-1119(92)90627-2.

34. Remmert, M., Biegert, A., Hauser, A., and Soding, J. (2011). HHblits: lightning-fast iterative protein sequence searching by HMM-HMM alignment. Nature methods 9, 173–175. 10.1038/nmeth.1818.

35. Mirdita, M., von den Driesch, L., Galiez, C., Martin, M.J., Soding, J., and Steinegger, M. (2017). Uniclust databases of clustered and deeply annotated protein sequences and alignments. Nucleic Acids Res 45, D170–D176. 10.1093/nar/gkw1081.

36. Steinegger, M., Mirdita, M., and Soding, J. (2019). Protein-level assembly increases protein sequence recovery from metagenomic samples manyfold. Nature methods 16, 603–606. 10.1038/s41592-019-0437-4.

37. Mirdita, M., Schutze, K., Moriwaki, Y., Heo, L., Ovchinnikov, S., and Steinegger, M. (2022). ColabFold: making protein folding accessible to all. Nature methods 19, 679–682. 10.1038/s41592-022-01488-1.

38. Baek, M., Anishchenko, I., Humphreys, I.R., Cong, Q., Baker, D., and DiMaio, F. (2023). Efficient and accurate prediction of protein structure using RoseTTAFold2. bioRxiv.

39. Anishchenko, I., Baek, M., Park, H., Hiranuma, N., Kim, D.E., Dauparas, J., Mansoor, S., Humphreys, I.R., and Baker, D. (2021). Protein tertiary structure prediction and refinement using deep learning and Rosetta in CASP14. Proteins 89, 1722–1733. 10.1002/prot.26194.

40. Kieser, T., Bibb, M.J., Buttner, M.J., Chater, K.F., and Hopwood, D.A. (2000). Practical Streptomyces Genetics (Crowes).

41. Ting, S.Y., LaCourse, K.D., Ledvina, H.E., Zhang, R., Radey, M.C., Kulasekara, H.D., Somavanshi, R., Bertolli, S.K., Gallagher, L.A., Kim, J., et al. (2022). Discovery of coordinately regulated pathways that provide innate protection against interbacterial antagonism. Elife 11. 10.7554/eLife.74658.

42. Tyanova, S., Temu, T., and Cox, J. (2016). The MaxQuant computational platform for mass spectrometry-based shotgun proteomics. Nature protocols 11, 2301–2319. 10.1038/nprot.2016.136.

43. Suloway, C., Pulokas, J., Fellmann, D., Cheng, A., Guerra, F., Quispe, J., Stagg, S., Potter, C.S., and Carragher, B. (2005). Automated molecular microscopy: the new Leginon system. J Struct Biol 151, 41–60. 10.1016/j.jsb.2005.03.010.

44. Rohou, A., and Grigorieff, N. (2015). CTFFIND4: Fast and accurate defocus estimation from electron micrographs. J Struct Biol 192, 216–221. 10.1016/j.jsb.2015.08.008.

45. Voss, N.R., Yoshioka, C.K., Radermacher, M., Potter, C.S., and Carragher, B. (2009). DoG Picker and TiltPicker: software tools to facilitate particle selection in single particle electron microscopy. J Struct Biol 166, 205–213.

46. Zivanov, J., Nakane, T., Forsberg, B.O., Kimanius, D., Hagen, W.J., Lindahl, E., and Scheres, S.H. (2018). New tools for automated high-resolution cryo-EM structure determination in RELION-3. Elife 7. 10.7554/eLife.42166.

47. Punjani, A., Rubinstein, J.L., Fleet, D.J., and Brubaker, M.A. (2017). cryoSPARC: algorithms for rapid unsupervised cryo-EM structure determination. Nature methods 14, 290–296. 10.1038/nmeth.4169.

48. Tegunov, D., and Cramer, P. (2019). Real-time cryo-electron microscopy data preprocessing with Warp. Nature methods 16, 1146–1152. 10.1038/s41592-019-0580-y.

49. Zivanov, J., Nakane, T., and Scheres, S.H.W. (2019). A Bayesian approach to beam-induced motion correction in cryo-EM single-particle analysis. IUCrJ 6, 5–17. 10.1107/S205225251801463X.

50. Rosenthal, P.B., and Henderson, R. (2003). Optimal determination of particle orientation, absolute hand, and contrast loss in single-particle electron cryomicroscopy. J Mol Biol 333, 721–745. 10.1016/j.jmb.2003.07.013.

51. Zhang, J., Tao, R., Liu, C., Wu, W., Zhang, Y., Cui, J., and Wang, J. (2013). Possible effects of iron deposition on the measurement of DTI metrics in deep gray matter nuclei: an in vitro and in vivo study. Neurosci Lett 551, 47–52. 10.1016/j.neulet.2013.07.003.

52. Wang, R.Y., Song, Y., Barad, B.A., Cheng, Y., Fraser, J.S., and DiMaio, F. (2016). Automated structure refinement of macromolecular assemblies from cryo-EM maps using Rosetta. Elife 5. 10.7554/eLife.17219.

53. de Moraes, M.H., Hsu, F., Huang, D., Bosch, D.E., Zeng, J., Radey, M.C., Simon, N., Ledvina, H.E., Frick, J.P., Wiggins, P.A., et al. (2021). An interbacterial DNA deaminase toxin directly mutagenizes surviving target populations. Elife 10. 10.7554/eLife.62967.

54. Gallagher, L.A., Velazquez, E., Peterson, S.B., Charity, J.C., Radey, M.C., Gebhardt, M.J., Hsu, F., Shull, L.M., Cutler, K.J., Macareno, K., et al. (2022). Genome-wide protein-DNA interaction site mapping in bacteria using a double-stranded DNA-specific cytosine deaminase. Nat Microbiol 7, 844–855. 10.1038/s41564-022-01133-9.

55. Gao, C., Montoya, L., Xu, L., Madera, M., Hollingsworth, J., Purdom, E., Singan, V., Vogel, J., Hutmacher, R.B., Dahlberg, J.A., et al. (2020). Fungal community assembly in drought-stressed sorghum shows stochasticity, selection, and universal ecological dynamics. Nat Commun 11, 34. 10.1038/s41467-019-13913-9.

56. Xu, L., Naylor, D., Dong, Z., Simmons, T., Pierroz, G., Hixson, K.K., Kim, Y.M., Zink, E.M., Engbrecht, K.M., Wang, Y., et al. (2018). Drought delays development of the sorghum root microbiome and enriches for monoderm bacteria. Proc Natl Acad Sci U S A 115, E4284–E4293. 10.1073/pnas.1717308115.

57. Schindelin, J., Arganda-Carreras, I., Frise, E., Kaynig, V., Longair, M., Pietzsch, T., Preibisch, S., Rueden, C., Saalfeld, S., Schmid, B., et al. (2012). Fiji: an open-source platform for biological-image analysis. Nature methods 9, 676–682. 10.1038/nmeth.2019.

58. Cutler, K.J., Stringer, C., Lo, T.W., Rappez, L., Stroustrup, N., Brook Peterson, S., Wiggins, P.A., and Mougous, J.D. (2022). Omnipose: a high-precision morphology-independent solution for bacterial cell segmentation. Nature methods 19, 1438–1448. 10.1038/s41592-022-01639-4.

59. Altschul, S.F., Madden, T.L., Schaffer, A.A., Zhang, J., Zhang, Z., Miller, W., and Lipman, D.J. (1997). Gapped BLAST and PSI-BLAST: a new generation of protein database search programs. Nucleic Acids Res 25, 3389–3402.

60. Aravind, L., Iyer, L.M., and Burroughs, A.M. (2022). Discovering Biological Conflict Systems Through Genome Analysis: Evolutionary Principles and Biochemical Novelty. Annu Rev Biomed Data Sci 5, 367–391. 10.1146/annurev-biodatasci-122220-101119.

61. Eddy, S.R. (2011). Accelerated Profile HMM Searches. PLoS computational biology 7, e1002195. 10.1371/journal.pcbi.1002195.

62. Mistry, J., Chuguransky, S., Williams, L., Qureshi, M., Salazar, G.A., Sonnhammer, E.L.L., Tosatto, S.C.E., Paladin, L., Raj, S., Richardson, L.J., et al. (2021). Pfam: The protein families database in 2021. Nucleic Acids Res 49, D412–D419. 10.1093/nar/gkaa913.

63. Kall, L., Krogh, A., and Sonnhammer, E.L. (2004). A combined transmembrane topology and signal peptide prediction method. J Mol Biol 338, 1027–1036. 10.1016/j.jmb.2004.03.016.

64. Lassmann, T. (2019). Kalign 3: multiple sequence alignment of large data sets. Bioinformatics 36, 1928–1929. 10.1093/bioinformatics/btz795.

65. Edgar, R.C. (2004). MUSCLE: multiple sequence alignment with high accuracy and high throughput. Nucleic Acids Res 32, 1792–1797.

66. Pei, J., and Grishin, N.V. (2014). PROMALS3D: multiple protein sequence alignment enhanced with evolutionary and three-dimensional structural information. Methods in molecular biology (Clifton, N.J 1079, 263–271. 10.1007/978-1-62703-646-7_17.

67. Taylor, W.R. (1986). The classification of amino acid conservation. J Theor Biol 119, 205–218. 10.1016/s0022-5193(86)80075-3.

68. Holm, L., and Rosenstrom, P. (2010). Dali server: conservation mapping in 3D. Nucleic Acids Res 38, W545–549. gkq366 [pii] 10.1093/nar/gkq366.

69. van Kempen, M., Kim, S.S., Tumescheit, C., Mirdita, M., Lee, J., Gilchrist, C.L.M., Soding, J., and Steinegger, M. (2023). Fast and accurate protein structure search with Foldseek. Nat Biotechnol. 10.1038/s41587-023-01773-0.

