## Supplementary material for "Umbrella toxin particles produced by Streptomyces block mycelial growth of competing species": Table S3

**Table S3 | Mass spectrometry-based identification of proteins immunoprecipitated by UmbC1, UmbC2, UmbC3 and UmbA1.** Data were filtered to remove proteins with fewer than 6 spectral counts in the IP samples for both biological replicates, and only those proteins enriched at least log 2 = 0.5 are shown, with the exception of UmbA-VSV-G, for which the most-enriched non-Umb protein was included for context.

| Bait protein | Protein ID | Control | IP | Enrichment (log2) |
|---|---|---|---|---|
| **UmbC1-VSV-G** | | | | |
| | UmbA4 | 0 | 49 | n/a |
| | UmbA5 | 0 | 40 | n/a |
| | UmbD1 | 0 | 28.5 | n/a |
| | UmbB1 | 0.5 | 18.5 | n/a |
| | UmbA6 | 0 | 17 | n/a |
| | UmbC1 | 1.5 | 134.5 | 6.02 |
| | UmbA1 | 6.5 | 53 | 2.91 |
| | Q8CJQ6 | 2 | 6.5 | 0.67 |
| | Q8CK05 | 6.5 | 15.5 | 0.73 |
| **UmbC2-VSV-G** | | | | |
| | UmbC2 | 0.5 | 46.5 | n/a |
| | UmbA4 | 0 | 12 | n/a |
| | Q9L0Y3 | 1 | 11.5 | n/a |
| | UmbA5 | 0.5 | 11 | n/a |
| | UmbB2 | 0 | 9.5 | n/a |
| | Q9KXP9 | 1.5 | 9.5 | n/a |
| | UmbC1 | 0.5 | 7.5 | n/a |
| | O54158 | 0.5 | 7 | n/a |
| | UmbA1 | 1 | 15.5 | 3.61 |
| | Q9RNU9 | 2.5 | 10.5 | 2.31 |
| | Q9S274 | 1.5 | 8.5 | 2.22 |
| | Q9XAC1 | 2 | 8 | 1.92 |
| | Q9XA42 | 2 | 7 | 1.80 |
| | Q9Z578 | 1.5 | 6 | 1.78 |
| | O69998 | 3 | 6 | 1.40 |
| | Q9FBR4 | 9.5 | 16.5 | 1.32 |
| | UmbA1 | 4 | 11 | 1.25 |
| | Q9RK35 | 3.5 | 9.5 | 1.25 |
| | Q9KZP4 | 2.5 | 7.5 | 1.24 |
| | Q9S2Q3\|Q9FC62 | 6.5 | 13.5 | 1.10 |
| | Q9L1Z5 | 3.5 | 8 | 1.08 |
| | Q9XAP2 | 2.5 | 6.5 | 1.08 |
| | Q9FC43 | 4 | 10 | 1.03 |

| | | | | |
|---|---|---|---|---|
| | Q9S2Q5\|Q9FC63 | 6 | 9.5 | 1.01 |
| | Q9XA23 | 5 | 7 | 1.01 |
| | Q7AKF3 | 8.5 | 15.5 | 0.98 |
| | Q9XAD1 | 7.5 | 18 | 0.94 |
| | Q9KXR6 | 8 | 15.5 | 0.93 |
| | Q9ZBU0 | 3.5 | 6.5 | 0.81 |
| | Q9Z564 | 6 | 13 | 0.76 |
| | Q9X909 | 8.5 | 14 | 0.67 |
| | Q9ADJ6 | 3.5 | 6 | 0.66 |
| | Q9EWV4 | 4.5 | 9 | 0.66 |
| | Q9RKK5 | 8 | 16 | 0.62 |
| | Q9L2H9 | 7.5 | 11 | 0.61 |
| | Q93RV9 | 4 | 7.5 | 0.60 |
| **UmbC3-VSV-G** | | | | |
| | UmbA3 | 0.5 | 61.5 | n/a |
| | UmbA4 | 0 | 45 | n/a |
| | UmbB3 | 0 | 32 | n/a |
| | UmbA6 | 0.5 | 26.5 | n/a |
| | UmbA2 | 0 | 9 | n/a |
| | UmbC3 | 10.5 | 781 | 4.71 |
| | UmbA5 | 2 | 62 | 3.79 |
| | UmbC1 | 1 | 9.5 | 1.70 |
| | O70007 | 1.5 | 9.5 | 1.30 |
| | Q9L244 | 2.5 | 8.5 | 0.92 |
| **UmbA1-VSV-G** | | | | |
| | UmbA4 | 0 | 51.5 | n/a |
| | UmbA5 | 0 | 49.5 | n/a |
| | UmbB1 | 0.5 | 38.5 | n/a |
| | UmbA6 | 0 | 26 | n/a |
| | UmbB2 | 0 | 11.5 | n/a |
| | UmbC1 | 1.5 | 147.5 | 5.99 |
| | UmbA1 | 6.5 | 145 | 4.41 |
| | Q9X8D4 | 5 | 8.5 | 0.23 |
