## Supplementary material for "Umbrella toxin particles produced by Streptomyces block mycelial growth of competing species": Table S5

### Table S5. Cryo-EM data collection, refinement and validation statistics

| | |
|---|---|
| **Data collection and processing** | |
| Magnification | |
| Voltage (kV) | 300 |
| Electron exposure (e–/Å$^2$) | 60 |
| Defocus range (μm) | -0.2 - -3.0 |
| Pixel size (Å) | 0.843 |
| Symmetry imposed | C1 |
| Initial particle images (no.) | 2,143,035 |
| Final particle images (no.) | 98,399 |
| Map resolution (Å) | 5.1 |
| FSC threshold | 0.143 |
| **Refinement** | |
| Initial model used (PDB code) | Alphafold-generated |
| Model resolution (Å) | 6.8 |
| FSC threshold | 0.5 |
| Model resolution range (Å) | 4.8-6.8 |
| Map sharpening *B* factor (Å$^2$) | -293 |
| Model composition | |
| Non-hydrogen atoms | 19,825 |
| Protein residues | 3,986 |
| Ligands | 0 |
| *B* factors (Å$^2$) | |
| Protein | 365.58 |
| Ligand | N/A |
| R.m.s. deviations | |
| Bond lengths (Å) | 0.009 |
| Bond angles (°) | 1.171 |
| Validation | |
| MolProbity score | 0.75 |
| Clashscore | 0.81 |
| Poor rotamers (%) | 0.00 |
| Ramachandran plot | |
| Favored (%) | 98.21 |
| Allowed (%) | 1.77 |
| Disallowed (%) | 0.03 |
