## Supplementary material for "Umbrella toxin particles produced by Streptomyces block mycelial growth of competing species": Table S6

**Table S6: Growth conditions for and results from screening Umb supernatant for growth inhibitory activity against diverse bacteria.** Growth was measured at a single timepoint using an ATP quantification-based bacterial cell viability assay. The average ratio represents the viability of each strain grown with control supernatant divided by Umb supernatant treatment. Z-scores were calculated from the average of log 2-transformed ratios from across all strains screened.

| Phylum | Genus and species | Source | Media | Average ratio | Z-score |
|---|---|---|---|---|---|
| Actinomycetota | *Allokutzneria albata* | NRRL No. B-24461 | ISP2 | 1.215 | 0.296 |
| Actinomycetota | *Amycolatopsis kentuckyensis* SAI_225 | This study | TSBY | 0.443 | -0.566 |
| Actinomycetota | *Amycolatopsis sp.* SAI_101 | This study | TSBY | 2.085 | 0.757 |
| Actinomycetota | *Arthrobacter nitroguajacolicus* SAI_060 | This study | TSBY | 0.667 | -0.217 |
| Actinomycetota | *Arthrobacter oxydans* SAI_217 | This study | TSBY | 0.516 | -0.436 |
| Actinomycetota | *Brevibacterium lyticum* | NRRL No. B-4262 | TGY | 0.781 | -0.081 |
| Actinomycetota | *Catenulispora acidophila* | NRRL No. B-24433 | ISP2 | 0.780 | -0.083 |
| Actinomycetota | *Cellulosimicrobium cellulans* | NRRL No. B-2381 | TGY | 0.503 | -0.457 |
| Actinomycetota | *Cellulosimicrobium cellulans* SAI_258 | This study | TSBY | 0.469 | -0.517 |
| Actinomycetota | *Cellulosimicrobium sp.* SAI_197 | This study | TSBY | 0.419 | -0.613 |
| Actinomycetota | *Dermacoccus sp.* SAI_028 | This study | TSBY | 1.222 | 0.301 |
| Actinomycetota | *Dermatophilus congolensis* | NRRL No. B-2350 | TSBY | 0.324 | -0.833 |
| Actinomycetota | *Frankia sp.* | NRRL No. B-16219 | ISP2 | 1.310 | 0.360 |
| Actinomycetota | *Georgenia sp.* | NRRL No. B-59275 | TGY | 0.484 | -0.491 |
| Actinomycetota | *Glycomyces rutgersensis* | NRRL No. B-16106 | ISP2 | 0.246 | -1.069 |
| Actinomycetota | *Kibdelosporangium aridum* | NRRL No. B-16436 | ISP2 | 1.098 | 0.210 |
| Actinomycetota | *Kineococcus gynurae* | NRRL No. B-24568 | ISP2 | 0.233 | -1.114 |
| Actinomycetota | *Kitasatospora kifunensis* | NRRL No. B-24284 | LB | 4.420 | 1.399 |
| Actinomycetota | *Kribbella hippodromi* | NRRL No. B-24553 | ISP2 | 0.855 | -0.004 |
| Actinomycetota | *Leifsonia sp.* 004_C5 | This study | TSBY | 1.406 | 0.420 |
| Actinomycetota | *Leifsonia sp.* 006_B1 | This study | TSBY | 0.891 | 0.031 |
| Actinomycetota | *Microbacterium atlanticum* SAI_030 | This study | TSBY | 0.794 | -0.067 |
| Actinomycetota | *Microbacterium jejuense* SAI_031 | This study | TSBY | 0.186 | -1.308 |
| Actinomycetota | *Microbacterium paraoxydans* SAI_221 | This study | TSBY | 0.370 | -0.720 |
| Actinomycetota | *Microbacterium trichothecenolyticum* CFG_272 | This study | TSBY | 0.613 | -0.288 |
| Actinomycetota | *Microtetraspora glauca* | NRRL No. B-3735 | ISP2 | 0.982 | 0.114 |
| Actinomycetota | *Mycobacterium goodii* 002_G6 | This study | TSBY | 1.979 | 0.713 |
| Actinomycetota | *Mycobacterium smegmatis* 001_A6 | This study | TSBY | 0.317 | -0.852 |
| Actinomycetota | *Nocardioides luteus* | NRRL No. B-16231 | R2A | 1.242 | 0.315 |
| Actinomycetota | *Nocardioides sp.* HB12 | PMID:29556109 | R2A | 0.854 | -0.006 |

| | | | | | |
|---|---|---|---|---|---|
| Actinomycetota | *Nocardioides sp.* SAI_065 | This study | TSBY | 0.748 | -0.119 |
| Actinomycetota | *Oerskovia turbata* | NRRL No. B-8019 | ISP2 | 1.000 | 0.129 |
| Actinomycetota | *Patulibacter minatonensis* | NRRL No. B-24346 | ISP2 | 0.575 | -0.343 |
| Actinomycetota | *Promicromonospora kroppenstedtii* TBS_116 | This study | TSBY | 1.139 | 0.241 |
| Actinomycetota | *S. albogriseolus* SAI_173 | This study | TSBY | 1.631 | 0.547 |
| Actinomycetota | *S. albogriseolus* SAI_190 | This study | TSBY | 0.865 | 0.005 |
| Actinomycetota | *S. ambofaciens* | NRRL No. B-2516 | TSBY | 0.645 | -0.245 |
| Actinomycetota | *S. ambofaciens* SAI_080 | This study | TSBY | 345.963 | 5.124 |
| Actinomycetota | *S. ambofaciens* SAI_104 | This study | TSBY | 14.312 | 2.403 |
| Actinomycetota | *S. ambofaciens* SAI_108 | This study | TSBY | 0.455 | -0.542 |
| Actinomycetota | *S. ambofaciens* SAI_195 | This study | TSBY | 1817.167 | 6.541 |
| Actinomycetota | *S. antibioticus* | NRRL No. B-1701 | TSBY | 0.465 | -0.524 |
| Actinomycetota | *S. anulatus* | NRRL No. B-2000 | TSBY | 0.924 | 0.062 |
| Actinomycetota | *S. azureus* | NRRL No. B-2655 | TSBY | 0.879 | 0.019 |
| Actinomycetota | *S. cellulosae* SAI_051 | This study | TSBY | 0.623 | -0.275 |
| Actinomycetota | *S. coelicolor* | NRRL No. B-3062 | TSBY | 0.888 | 0.029 |
| Actinomycetota | *S. collinus* SAI_078 | This study | TSBY | 8.157 | 1.923 |
| Actinomycetota | *S. corchorusii* SAI_180 | This study | TSBY | 1.269 | 0.333 |
| Actinomycetota | *S. eurocidicus* | NRRL No. B-1701 | TSBY | 1.219 | 0.299 |
| Actinomycetota | *S. graminofaciens* SAI_110 | This study | TSBY | 0.914 | 0.052 |
| Actinomycetota | *S. graminofaciens* SAI_175 | This study | TSBY | 0.795 | -0.067 |
| Actinomycetota | *S. graminofaciens* SAI_211 | This study | TSBY | 1.065 | 0.184 |
| Actinomycetota | *S. griseorubiginosus* SAI_142 | This study | TSBY | 0.863 | 0.004 |
| Actinomycetota | *S. griseus* | NRRL No. B-2682 | TSBY | 25.362 | 2.892 |
| Actinomycetota | *S. lienomycini* SAI_166 | This study | TSBY | 32.319 | 3.099 |
| Actinomycetota | *S. lividans* | NRRL No. B65306 | TSBY | 0.969 | 0.103 |
| Actinomycetota | *S. luteogriseus* | NRRL No. B-12422 | TSBY | 6.490 | 1.727 |
| Actinomycetota | *S. mobaraensis* | NRRL No. B-3729 | TSBY | 1.282 | 0.342 |
| Actinomycetota | *S. pristinaespiralis* | NRRL No. B2958 | TSBY | 0.993 | 0.124 |
| Actinomycetota | *S. pristinaespiralis* SAI_178 | This study | TSBY | 0.632 | -0.263 |
| Actinomycetota | *S. rochei* SAI_164 | This study | TSBY | 4.238 | 1.363 |
| Actinomycetota | *S. sp.* SAI_041 | This study | TSBY | 0.672 | -0.210 |
| Actinomycetota | *S. sp.* SAI_103 | This study | TSBY | 5.954 | 1.654 |
| Actinomycetota | *S. sp.* SAI_117 | This study | TSBY | 0.833 | -0.027 |
| Actinomycetota | *S. sp.* SAI_126 | This study | TSBY | 0.848 | -0.011 |
| Actinomycetota | *S. sp.* SAI_133 | This study | TSBY | 0.950 | 0.085 |
| Actinomycetota | *S. sp.* SAI_167 | This study | TSBY | 3.824 | 1.275 |
| Actinomycetota | *S. sp.* SAI_203 | This study | TSBY | 0.788 | -0.074 |

| | | | | | |
|---|---|---|---|---|---|
| Actinomycetota | *S. tendae* SAI_182 | This study | TSBY | 0.415 | -0.621 |
| Actinomycetota | *S. tendae* SAI_185 | This study | TSBY | 0.827 | -0.033 |
| Actinomycetota | *S. tendae* SAI_218 | This study | TSBY | 0.809 | -0.051 |
| Actinomycetota | *Saccharothrix coeruleofusca* | NRRL No. B-16115 | LB | 5.768 | 1.626 |
| Actinomycetota | *Sanguibacter sp.* | NRRL No. B-59339 | TGY | 0.587 | -0.326 |
| Actinomycetota | *Terrabacter tumescens* | NRRL No. B-4012 | TGY | 0.501 | -0.461 |
| Bacteroidota | *Chryseobacterium bacterium* CI02 | PMID:16885294 | TSBY | 0.664 | -0.220 |
| Bacteroidota | *Flavobacterium johnsoniae* UW101 | Gift from Mark McBride's lab | LB | 0.797 | -0.065 |
| Bacteroidota | *Sphingobacterium bacterium* CI01 | PMID:16885294 | TSBY | 0.753 | -0.112 |
| Firmicutes | *Alkalihalobacillus alcalophilus* | NRRL No. B-14309 | Alkaline Nutrient Agar | 1.003 | 0.132 |
| Firmicutes | *Bacillus aryabhattai* TBS_067 | This study | TSBY | 0.496 | -0.469 |
| Firmicutes | *Bacillus haynesii* TBS_113 | This study | TSBY | 0.566 | -0.356 |
| Firmicutes | *Bacillus megaterium* 005_H9 | This study | TSBY | 0.605 | -0.300 |
| Firmicutes | *Bacillus pumilus* TBS_051 | This study | TSBY | 0.502 | -0.459 |
| Firmicutes | *Bacillus pumilus* TBS_099 | This study | TSBY | 0.546 | -0.388 |
| Firmicutes | *Bacillus sp.* 005_H12 | This study | TSBY | 0.541 | -0.396 |
| Firmicutes | *Bacillus sp.* 006_D12 | This study | TSBY | 0.485 | -0.489 |
| Firmicutes | *Bacillus subtilis* 004_F4 | This study | TSBY | 0.635 | -0.258 |
| Firmicutes | *Bacillus subtilis* 006_C4 | This study | TSBY | 0.501 | -0.462 |
| Firmicutes | *Bacillus velezensis* 006_A12 | This study | TSBY | 0.661 | -0.224 |
| Firmicutes | *Bacillus zanthoxyli* TBS_040 | This study | TSBY | 2.024 | 0.732 |
| Firmicutes | *Bacillus zanthoxyli* TBS_056 | This study | TSBY | 0.573 | -0.346 |
| Firmicutes | *Brevibacillus borstelensis* | NRRL No. NRS-948 | TGY | 0.395 | -0.663 |
| Firmicutes | *Brevibacillus chosinensis* | NRRL No. B-23247 | TGY | 0.528 | -0.417 |
| Firmicutes | *Brevibacillus laterosporus* | NRRL No. NRS-1339 | TGY | 0.401 | -0.650 |
| Firmicutes | *Brevibacillus parabrevis* | NRRL No. NRS-751 | TGY | 0.459 | -0.536 |
| Firmicutes | *Carnobacterium divergens* | NRRL No. B-23835 | Liver Infusion Broth | 0.555 | -0.374 |
| Firmicutes | *Carnobacterium gallinarum* | NRRL No. B-14832 | Liver Infusion Broth | 0.408 | -0.636 |
| Firmicutes | *Carnobacterium maltaromaticum* | NRRL No. B-14829 | Liver Infusion Broth | 0.368 | -0.724 |
| Firmicutes | *Caryophanon latum* | NRRL No. B-1893 | ISP2 | 0.544 | -0.390 |
| Firmicutes | *Cytobacillus kochii* | NRRL No. NRS-1758 | TGY | 0.335 | -0.805 |
| Firmicutes | *Cytobacillus praedii* | NRRL No. B-14566 | TGY | 0.450 | -0.552 |
| Firmicutes | *Fictibacillus marinisediminis* | NRRL No. B-59209 | TGY | 1.065 | 0.184 |
| Firmicutes | *Gracibacillus dipsosauri* | NRRL No. B-23348 | TSBY | 0.588 | -0.323 |
| Firmicutes | *Heyndrickxia sporothermodurnas* | NRRL No. NRS-1638 | TGY | 0.547 | -0.387 |
| Firmicutes | *Kurthia gibsonii* | NRRL No. B-41085 | TGY | 0.210 | -1.205 |
| Firmicutes | *Kurthia sibirica* | NRRL No. B-41083 | TGY | 0.620 | -0.279 |

| | | | | | |
|---|---|---|---|---|---|
| Firmicutes | *Lactococcus lactis* | NRRL No. B-23804 | Liver Infusion Broth | 0.295 | -0.914 |
| Firmicutes | *Leuconostoc citreum* | NRRL No. B-1501 | Liver Infusion Broth | 0.455 | -0.544 |
| Firmicutes | *Lysinibacillus fusiformis* | NRRL No. B-14865 | TGY | 0.460 | -0.534 |
| Firmicutes | *Lysinibacillus xylanityticus* | NRRL No. NRS-1307 | TGY | 0.487 | -0.485 |
| Firmicutes | *Neobacillus mesonae* | NRRL No. B-14565 | TGY | 0.591 | -0.320 |
| Firmicutes | *Oenococcus oeni* | NRRL No. B-3474 | Liver Infusion Broth | 0.514 | -0.439 |
| Firmicutes | *Paenibacillus alba* | NRRL No. BD-533 | TGY | 0.898 | 0.038 |
| Firmicutes | *Paenibacillus amylolyticus* | NRRL No. B-14940 | TGY | 1.267 | 0.332 |
| Firmicutes | *Paenibacillus chibensis* | NRRL No. B-142 | TGY | 0.388 | -0.680 |
| Firmicutes | *Paenibacillus favisporus* 004_C2 | This study | TSBY | 0.599 | -0.308 |
| Firmicutes | *Paenibacillus lautus* 004_C1 | This study | TSBY | 0.568 | -0.354 |
| Firmicutes | *Paenibacillus lautus* TBS_091 | This study | TSBY | 1.048 | 0.169 |
| Firmicutes | *Paenibacillus lautus* TBS_092 | This study | TSBY | 0.238 | -1.096 |
| Firmicutes | *Pediococcus pentosaceus* | NRRL No. B-14620 | Liver Infusion Broth | 0.394 | -0.666 |
| Firmicutes | *Peribacillus castrilensis* | NRRL No. B-41276 | TGY | 0.281 | -0.955 |
| Firmicutes | *Peribacillus frigoritolerans* | NRRL No. BD-432 | TGY | 0.571 | -0.349 |
| Firmicutes | *Peribacillus simplex* | NRRL No. BD-267 | TGY | 0.219 | -1.167 |
| Firmicutes | *Planococcus sierraensis* | NRRL No. B-65582 | R2A | 0.620 | -0.279 |
| Firmicutes | *Priestia aryabhattai* | NRRL No. BD-263 | TGY | 1.628 | 0.546 |
| Firmicutes | *Priestia endophytica* | NRRL No. BD-290 | TGY | 0.631 | -0.264 |
| Firmicutes | *Rummeliibacillus stabekisii* | NRRL No. B-51320 | TSBY | 0.441 | -0.569 |
| Firmicutes | *Sporosarcina globispora* | NRRL No. B-3396 | TGY | 0.827 | -0.032 |
| Firmicutes | *Staphylococcus sp.* HA57 | PMID:29556109 | R2A | 0.575 | -0.343 |
| Firmicutes | *Sutcliffiella cohnii* | NRRL No. B-14735 | Alkaline Nutrient Agar | 0.465 | -0.524 |
| Firmicutes | *Weissella confusa* | NRRL No. B-1064 | Liver Infusion Broth | 0.470 | -0.515 |
| Firmicutes | *Weissella viridescens* | NRRL No. B-1951 | Liver Infusion Broth | 0.360 | -0.742 |
| Proteobacteria | *Acinetobacter baylyi* ADP1 | ATCC #33305 | BHI | 0.640 | -0.252 |
| Proteobacteria | *Agrobacterium tumefaciens* FACH | Dong *et al.* 1992. *Phytopathology.* | TSBY | 0.810 | -0.051 |
| Proteobacteria | *Burkholderia thailandensis* E264 | PMID:16725056 | LB | 1.129 | 0.233 |
| Proteobacteria | *Erwinia caratovora* Ecc71 | PMID:9701816 | LB | 0.950 | 0.086 |
| Proteobacteria | *Escherichia coli* MG1655 | ATCC #47076 | LB | 0.498 | -0.466 |
| Proteobacteria | *Lysobacter enzymogenes* UASM495 | ATCC #29487 | TSBY | 0.928 | 0.065 |
| Proteobacteria | *Pseudomonas aeruginosa* PAO1 | PMID:10984043 | LB | 0.736 | -0.132 |
| Proteobacteria | *Serratia proteamaculans* | PMID:26187596 | LB | 0.838 | -0.021 |
| Proteobacteria | *Xanthomonas maltophilia* | ATCC #13637 | TSBY | 0.669 | -0.213 |
